# Genome-wide association study of morphometric and metabolic characteristics in the European populations of the sugar kelp *Saccharina latissima*

**DOI:** 10.64898/2026.04.27.720995

**Authors:** Stéphane Mauger, Komlan Avia, Lucie Jaugeon, Paolo Ruggeri, Zofia Nehr, Ousseini Issaka Salia, Jérôme Coudret, Emilie Gouhier, Aurélien Baud, Stéphane Loisel, Antoine Fort, Ronan Sulpice, Christophe Destombe, Philippe Potin, J. Mark Cock, Myriam Valero

## Abstract

The sugar kelp *Saccharina latissima* is a promising candidate for sustainable aquaculture in the North Atlantic and North-East Pacific but genetic improvement has been hindered by limited understanding of the genetic basis of economically important traits. We conducted the first genome-wide association study (GWAS) for this species using 202 self-fertilised pseudo-F1 individuals derived from 12 populations spanning northern and southern European genetic clusters. Individuals were genotyped with ddRAD-seq-derived SNP markers and phenotyped in a common garden experiment for four morphological traits (blade length, blade width, blade area, stipe length) and six metabolic traits related to nitrogen metabolism. We identified 26 significant marker-trait associations, with phenotypic variance explained (PVE) ranging from 0.65% to 52.44%. Major-effect loci were detected for blade width (52.44% PVE) and blade area (45.22% PVE) and a locus on chromosome 17 influenced both blade length and blade area. Marker-based heritability estimates ranged from 0.75 to 0.99 for morphological traits and from 0.00 to 0.99 for metabolic traits, though with large standard errors. Cross-validation of genomic selection models yielded predictive abilities of 0.21-0.59 across traits. Our findings reveal a mixed genetic architecture with major-effect loci suitable for marker-assisted selection and polygenic traits amenable to genomic selection, providing a foundation for genomics-assisted breeding programs in kelp aquaculture.

## Introduction

The sugar kelp *Saccharina latissima* (Linnaeus) (Lane et al., 2006) is an ecologically and economically important species. It is a sister species of the Asian *Saccharina japonica* (J.E. Areschoug) (Lane et al., 2006) which represents a global production of 10-12 millions fresh tons (FAO, 2024), and it is exploited commercially in Europe and North America, although at a scale an order of magnitude lower than reported for *S. japonica*. *S. latissima* is used for human consumption, animal feed, and as a potential source of diverse bioactive compounds (e.g. alginate, fucoidans, glucans and iodine) that have applications in a wide range of industrial processes (see review by Sæther et al. (2024). Owing to its high growth rate and broad environmental tolerance, *S. latissima* is used for commercial cultivation, either as a monoculture or within integrated multi-trophic aquaculture (IMTA) systems (Broch et al., 2019; Mildenberger et al., 2022). Several countries, particularly in Europe, have invested in the development of *S. latissima* aquaculture. As seaweed farming is a rapidly expanding sector, there is an urgent need to develop efficient and sustainable breeding strategies to improve biomass yield and quality while minimizing environmental impacts.

Wild populations of *S. latissima* have a broad geographic distribution, spanning the northern hemisphere from polar to temperate regions. Neiva and collaborators (2018) identified four distinct phylogroups, based on mitochondrial COI and nuclear SSR molecular markers: the first detected in the NE and NW Pacific, W Greenland and the Hudson Bay, the second in the NW Atlantic, the third in the NE Atlantic (European phylogroup) and the fourth restricted to Russia’s Pacific region of Primorye. In the European phylogroup, the pattern of genetic structure has been studied using on COI, SSR and SNP markers among distant populations sampled from Spitsbergen (Svalbard archipelago, Eastern Greenland Sea) to Portugal (Diehl et al., 2023; Guzinski et al., 2020; Jaugeon et al., 2025; Neiva et al., 2018). Mitochondrial and nuclear molecular markers were all congruent in these papers, indicating strong genetic differentiation between northern and southern Europe. Moreover, Jaugeon et al., (2025) further demonstrated that southern and northern European populations corresponded to distinct genetic clusters that arose from historical processes implying recolonization after the Last Glacial Maximum and originating from various refugia. Furthermore, the Jaugeon et al., (2025) study helped to better define the boundaries between the two clusters: the southern boundary of the northern cluster extending to southern Ireland, and the northern boundary of the southern cluster to the English Channel and the southern part of the North Sea. Higher genetic diversity and allelic richness was revealed in the northern cluster while higher genetic sub-structure was characteristic of the southern one.

The strong genetic structure of *S. latissima* can be largely attributed to its biological characteristics and limited dispersal capacity. This brown alga exhibits a heteromorphic haplo-diplontic life cycle and a short generation time spanning 1 to 2 years. Dispersal is restricted to short-lived mobile stages (spores and gametes), whose transport relies primarily on ocean currents, thereby constraining gene flow among populations (Ribeiro et al., 2022).

Kelp species are well known for their pronounced morphological plasticity and high adaptive potential (Bartsch et al., 2008). Although a recent large-scale study of adult *S. latissima* across a broad latitudinal gradient in the NE Atlantic reported weak correlations between morphological and biochemical traits and abiotic environmental parameters (Diehl et al., 2023), experimental evidence clearly demonstrates that temperature and light strongly influence thallus morphology, physiology, and growth. These effects have been consistently observed in common-garden experiments and controlled laboratory studies (Gerard, 1988; Gerard & Du Bois, 1988; Lüning et al., 1978). Importantly, these experimental studies also revealed population-specific responses to environmental conditions, providing strong evidence for local adaptation and ecotypic differentiation within *S. latissima* (Gerard, 1988; Gerard & Du Bois, 1988; Lüning et al., 1978; Umanzor et al., 2021). Although the genetic basis of adaptations remains relatively poorly understood, some SNP markers have been associated with brackish water environments (Møller Nielsen et al., 2016) and with temperature gradients (Guzinski et al., 2020), providing molecular evidence consistent with local adaptation. Based on this knowledge, recent breeding programs have been attempted at different geographical scales. For example, at the local level, using individuals from the NW Atlantic (Umanzor et al., 2021), or at the global level, using a large sample covering the worldwide distribution area of *S. latissima* (Cohen et al., 2025). These different approaches integrated morphological and physiological characteristics into genetic selection frameworks, with the dual objective of improving kelp cultivation performance and setting the framework for domestication of this species. Understanding the genetic diversity of local populations should facilitate the selection of strains essential for the development of seaweed aquaculture (Goecke et al., 2020).

Traditionally, breeding programs have identified quantitative trait loci (QTLs) by analyzing trait-marker co-segregation in biparental populations. In haplo-diplobiontic species such as kelp, however, the generation of suitable segregating populations for classical QTL mapping is particularly challenging (Nehr et al., 2025). Two recent QTL studies in *S. latissima* have nevertheless demonstrated exploitable genetic variance for traits related to temperature stress responses (Nehr et al., 2025) and key morphological traits associated with sporophyte size, including stipe length, blade width and blade length (Bråtelund et al., 2026). Notably, the latter study also reported substantial inbreeding depression, highlighting a major limitation of breeding strategies that implement crosses that involve at least some degree of selfing. Taken together, these findings indicate that kelp breeding programs would benefit substantially from genomics-based approaches such as genome-wide association studies (GWAS) and genomic selection (GS), which can leverage natural genetic variation while reducing the need for highly structured breeding populations and thereby accelerating genetic improvement.

Beyond QTL mapping, modern genomics-based approaches for crop improvement, such as GWAS and GS, are increasingly applied for crop improvement (Heslot et al., 2015). GWAS have revolutionized plant and animal breeding by enabling the identification of genetic variants associated with complex traits without requiring controlled crosses, instead exploiting natural variation within diverse germplasm collections (Korte & Farlow, 2013). By scanning thousands to millions of markers across the genome, GWAS can pinpoint candidate loci underlying traits of agronomic importance, from yield to stress tolerance (Huang & Han, 2014). Complementing GWAS, GS uses genome-wide marker information to predict breeding values, allowing selection of superior individuals at early developmental stages and significantly shortening breeding cycles (Crossa et al., 2017; Meuwissen et al., 2001). Importantly, GS overcomes key limitations of marker-assisted selection (MAS), which is effective only for traits controlled by few major loci but performs poorly for complex polygenic traits where many small-effect variants collectively determine phenotypic variation (Bernardo & Yu, 2007; Heslot et al., 2015). These approaches have proven particularly valuable in species with long generation times or complex life cycles, where traditional pedigree-based methods are inefficient (Grattapaglia & Resende, 2011). In algae, the application of GWAS and GS remains in its infancy but holds considerable promise. Recent advances in kelp genomics, including chromosome-level genome assemblies for *S. japonica* (Ye et al., 2015) and *S. latissima* (Denoeud et al., 2024; https://phaeoexplorer.sb-roscoff.fr/), now provide the foundational tools necessary to implement these powerful approaches for accelerating genetic gain in seaweed aquaculture.

In this study, we used a GWAS approach to identify genetic loci associated with economically important traits in the kelp species *S. latissima*. Our study focused on 13 populations sampled across the European distribution range, representing the northern and southern genetic clusters that reflect the species’ postglacial recolonization history. Self-fertilised pseudo-F1 progeny obtained from wild-collected sporophytes were cultivated in a common garden experiment and phenotyped for four morphological traits (blade length, blade width, blade area, stipe length), as proxies for growth and biomass production, as well as six metabolic traits related to nitrogen metabolism (NO₂, NO₃^-^, NH₄^+^, proteins, amino acids, and rhamnose) relevant to nutritional quality and cultivation performance. Kelps are potential sources of protein for animal feed and human food (Aasen et al., 2022; Marinho et al., 2015). They can potentially also be exploited for bio-remediation of nitrogen in marine ecosystems (Grebe et al., 2021; Roleda & Hurd, 2019). Our objectives were to: (1) dissect the genetic architecture of those key morphological and metabolic traits; (2) identify candidate genes and functional pathways underlying trait variation; (3) compare genetic patterns between northern and southern populations; and (4) assess the feasibility of marker-assisted selection and genomic selection as tools for kelp breeding programs. This study provides the first comprehensive GWAS in *S. latissima* and establishes a foundation for genomics-assisted breeding in this emerging aquaculture species.

## Materials and methods

### Sampling sites of the parental populations

Thirteen sites of *S. latissima* were selected across the European distribution of the species (Figure 1, Supplementary Table 1) from Spain to Sweden. A total of 365 parental sporophytes were collected with 25 to 38 individuals per site, except in LOC and PTW, where only 9 and 10 individuals could be found (Figure 2, Supplementary Table 1).

**Figure 1.**
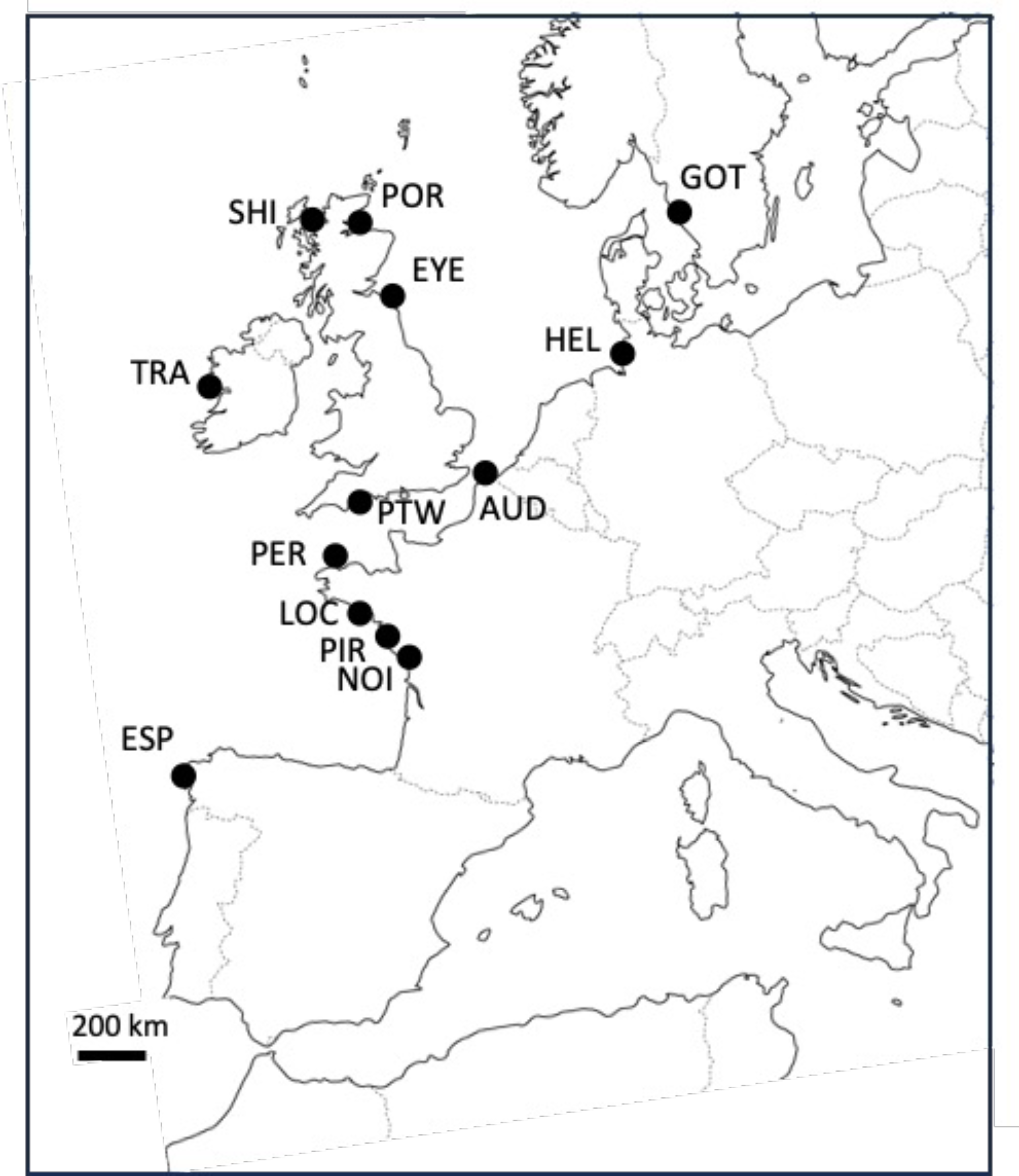
Location of the sampled sites. See Supplementary Table 1 for explanations of the location codes.

**Figure 2.**
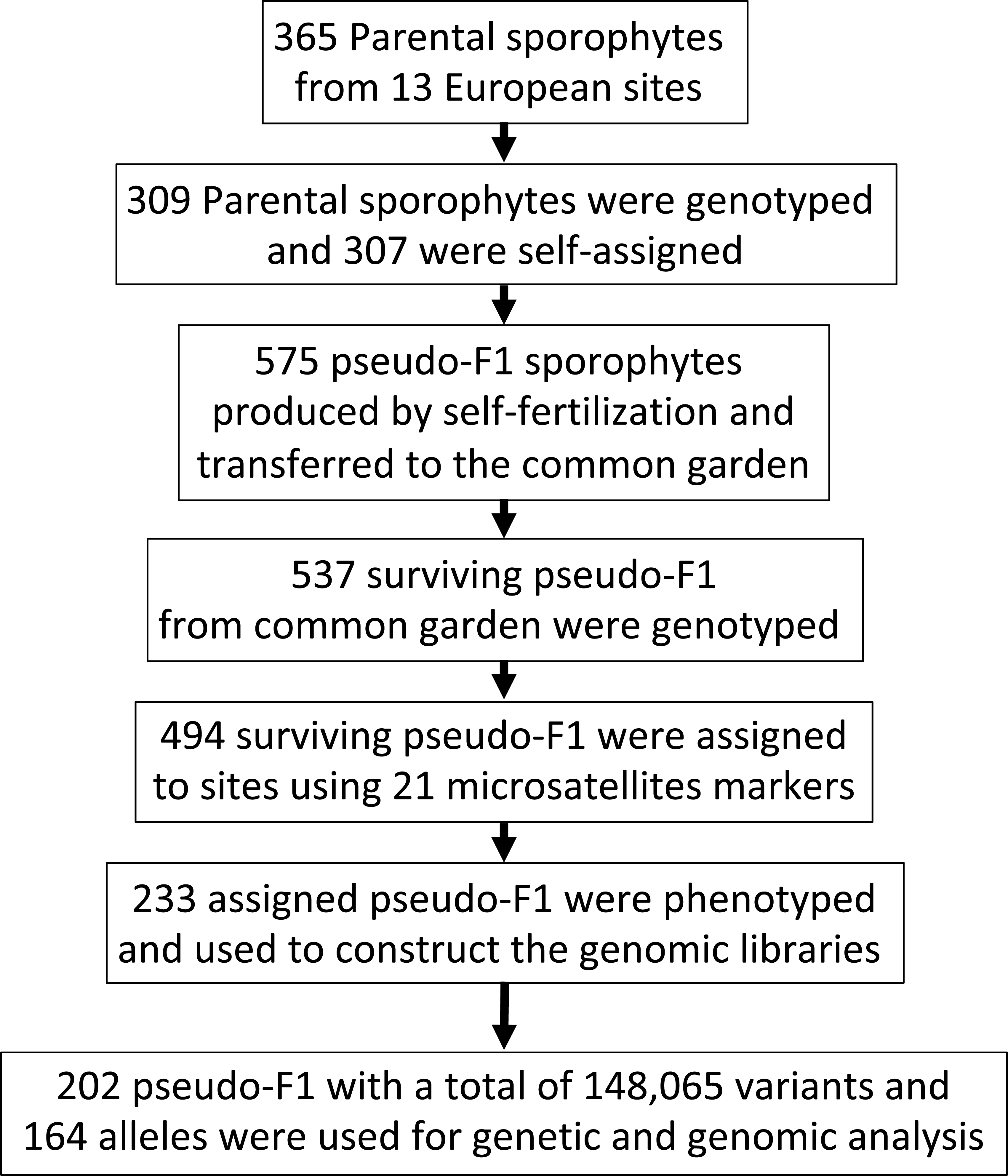
Workflow and experimental setup.

### Tissue collection and spore release

For each sporophyte, the blade was cleaned to remove all epiphytes, rinsed with sterile seawater and thoroughly blotted dry using paper towels. A 2 x 2 cm piece of vegetative blade tissue was taken in the meristematic area and stored with silica gel for future DNA extraction. Another 2 x 2 cm piece of fertile blade tissue was taken in the darkest part of the sorus and placed in a 50 mL falcon tube containing sterile seawater and two microscope glass slides for spore release. When the sporophyte was not mature, the sori were induced as described in Forbord et al., (2020). Then, the two microscope glass slides with the settled spores were transferred to 90 x 16 mm Petri dishes containing filtered (5 µm) sterile seawater enriched with half-strength Provasoli Enriched Seawater (herein referred to as P.E.S; Provasoli, 1968). The spores were allowed to develop to 5-10 cell gametophytes under vegetative growth culture conditions (13°C, red light, irradiance between 10-20 µmol.m^-^².s^-1^, 12:12 h light:dark photoperiod).

### Production of sporophytes from self-fertilized gametophytes: pseudo-F1 populations

Each pseudo-F1 individual was generated by crossing (i.e., self-fertilizing a pool of gametophytes derived from a single F0 sporophyte parent via multiple meiotic events). Self-fertilization of the gametophytes was carried out by transferring one of the microscope glass slides to a 145 x 20 mm Petri dish filled with half-strength P.E.S and placed at 13°C under white light with a light intensity between 20-30 µmol.m^-2^.s^-1^ and a light/dark cycle of 8/16 h during the first 4 days and 12/12 h photoperiod until the end of the experiment. The medium was changed every fortnight and the development of the young sporophytes was controlled at these times under an inverted microscope (Olympus CKX41/CKX31, Japan). After 30 days of culture, the young sporophytes measured approximately one millimeter and the microscope glass slide was transferred to a new 145 x 20 mm Petri dish. The slide was gently scraped with a cell scraper to detach the sporophytes and obtain a free-living culture. After 45 days of culture, the sporophytes measured a few millimeters and developed as “bouquets” (clusters of sporophytes). About ten of these “bouquets” were selected and transferred to a new 145 x 20 mm Petri dish. After 60 days of culture, the young sporophytes reached a length of approximately one centimeter and it was possible to separate them from each other using dissecting forceps and scalpel under a binocular microscope (Olympus SZ61, Japan).

### Common garden experiment

Once separated, the sporophytes were transferred to a common garden (Supplementary Figure 1). The experimental setup consisted of 10 L Nalgene flasks on shelves in a climate room. The cultures were kept at 13°C under white light with a light intensity between 30-40 µmol.m^-2^.s^-1^ at the surface of the flask. The light/dark cycle of 12/12h remained unchanged during the entire experiment. Light was provided by a fluorescent tube (Philips Master TL-D 18W/865). Filtered air (0.2 µm PTFE membrane, Whatman) was provided through silicon tubes attached to a plastic pipette for aeration. The location of the culture flasks on the shelves was changed randomly each time the medium was renewed. The seawater used during the common garden experiments was collected one month before the beginning of the cultures in flasks and stored in three 650 L tanks to ensure the same composition and quality of the seawater during the entire experiment.

Each culture flask contained sporophytes that originated from different populations (575 individuals in 30 culture flasks in total, see Supplementary Table 2, Figure 2). At the end of the experiment, the Pseudo-F1 sporophytes were assigned to their population of origin by running an assignment test based on multilocus genotypes using GenClass2 version 2.0.g (Piry et al., 2004) with an assignation threshold score of 0.05. The two first weeks (day 61 to 75), the flasks were filled with 5 L of filtered (1 µm), non-autoclaved seawater enriched with half-strength P.E.S. Then, the culture medium was renewed weekly with 8 L of filtered (1 µm), non-autoclaved seawater enriched with half-strength P.E.S the first time and enriched with full strength P.E.S the following times.

All cultures were harvested when the sporophytes reached a sufficient length to have enough material for analysis (approximately 90 days after the beginning of the self-fertilization of the gametophytes).

#### Phenotyping the pseudo-F1 populations

Harvested sporophytes were photographed (Supplementary Figure 2) with a backlight and a Nikon camera. These pictures were then analysed with ImageJ^®^ (Schneider et al., 2012) software in order to measure four different morphometric parameters: stipe length (cm), blade length (cm), blade width on the wider part of the frond (cm) and blade surface area (cm^2^). The meristematic area was cut and stored with silica gel for further DNA extraction. The remaining parts of the blade were put into a 2 mL Eppendorf tube, directly plunged in liquid nitrogen and then stored at -80°C. These tissues were then lyophilized, ground to powder and approximately 5 mg were used to measure the six metabolic traits: concentration of nitrite (NO_2_: µgN/gDW), nitrate (NO_3_^-^: µgN/gDW) protein (mg/gDW), amino acids (µmol/gDW), ammonium (NH_4_^+^: µmole/gDW), and rhamnose (µmole/gDW). Soluble metabolites were extracted with hot ethanolic extraction, and the insoluble pellet was resuspended in 0.1 M sodium hydroxide. The redissolved insoluble pellet was used to determine protein content using the Lowry method. The other assays used the hot ethanolic extracts. NO_2_ and NO_3_^-^ were determined using the Griess method, amino acids using ninhydrin, and rhamnose using the oxidation of NAD^+^ into NADH by a rhamnose-dehydrogenase enzyme (Megazyme, Ireland).

### Statistical analyses of morphological and metabolic traits

To evaluate the significance of differences between the geographical sites of origin for the five morphological and the six metabolic traits, we fitted a general linear model using Minitab® software (version: 19.2020.2.0): site of origin was considered as a fixed factor and individuals within sites as replicates of the trait measured for the given site. We only retained the sites in which the traits were measured in a minimum of five individuals (i.e., replicates per sites). Means between sites were compared using Tukey pairwise comparisons with Minitab® software. Mean morphological and metabolic traits were compared among genetic clusters using Kruskal-Wallis tests with Minitab® software. Correlation between traits was estimated using the function “chart.Correlation” in the R package “PerformanceAnalytics” version 2.0.8.

### DNA extraction, genotyping of SSR markers and assignment test of the pseudo-F1s to their parental populations

Total genomic DNA was extracted from approximately 10-20 mg of silica gel-dried tissue. DNA extraction was performed using the Nucleospin 96 plant kit (Macherey-Nagel, Germany) according to the manufacturer’s instructions, except that samples were left in PL1 lysis buffer at 65°C for 15 min and the washing steps with buffer PW1 and PW2 were each repeated twice. These modifications were performed to remove most of the polysaccharides that might inhibit DNA extraction and PCR amplification. DNA was eluted twice with 60 µL of PE buffer pre-heated to 70°C. The genomic DNA was purified using the Nucleospin gDNA Clean-up XS, (Macherey-Nagel, Germany) and purified according to the manufacturer’s instruction with final elution into 30 µL (2 x 15 µL) of supplied elution buffer.

Amplification and scoring of Simple Sequence Repeat (SSR) loci were performed as detailed in Guzinski et al. (2016) for 19 Expressed Sequences Tag (EST)-derived SSR loci, and as in Paulino et al. (2016) for 8 genomic SSR loci. Amplification products were separated by electrophoresis on an ABI 3130 XL capillary sequencer (Applied Biosystems, USA). Alleles were sized using the SM594 size standard (Mauger et al., 2012) and scored manually using the software GeneMapper 4.0 (Applied Biosystems). Two filters were applied to the raw multilocus genotype dataset using the poppr R package version 2.9.6 (Kamvar et al., 2014) using the parameters as detailed in Jaugeon et al. (2025). The final dataset included 309 of 365 parental sporophytes (Supplementary Table 3) and 537 of 575 pseudo-F1 sporophytes from the common garden experiment (Fig 2, Supplementary Table 2) and all individuals were characterized with 21 SSR loci, i.e. all 18 loci used in Guzinski et al. (2020) except Sacl-32 and Sacl-75, and 5 additional markers: Sacl-11/37/57/65/95 from Guzinski et al. (2016).

Assignment of individuals to their population of origin was performed using a method based on multilocus genotypes in GenClass2 version 2.0.g (Piry et al., 2004). The method was first validated by self-assigning parental individuals to their site of origin. The “assign/exclude populations as origin of individuals” option was employed for individual assignment, applying a threshold score of 0.05. Both the Bayesian method (Rannala & Mountain, 1997) and the frequency-based method (Paetkau et al., 2004) were used for assignment. Assignment results were validated only when both methods yielded the same assignment with a score of 90% or higher.

Using 21 SSR loci, this approach achieved an assignment success rate exceeding 99% for parental individuals to their population of origin (see Results, Supplementary Table 3). Consequently, we applied this method to assign flask-cultured pseudo-F1 individuals to their parental populations of origin.

### Construction of genomic libraries for ddRAD-seq, filtering and SNP discovery

We retained only individuals that could be both accurately assigned to their population of origin and phenotyped, resulting in a total of 233 out of 494 of the total viable pseudo-F1s individuals (see results, Supplementary Table 2, Figure 2), which were then used to construct the genomic libraries. The protocol for preparation of double digest restriction-site associated DNA (ddRAD) libraries (Mauger & Avia, 2024) was adapted from Peterson et al. (2012) and detailed for *S. latissima* in Nehr et al. (2025). The ddRAD library was sequenced (150 bp paired-end reads) on one lane of an Illumina NovaSeq6000 S4 platform to produce 4.281 billion raw reads. Quality control of the sequences was performed using FASTQC version 0.11.7 (Andrews et al., 2010). Adaptors were removed, and the reads were trimmed to 138 bp using Trimmomatic version 0.39 (Bolger et al., 2014). After demultiplexing, filtering of low quality and ambiguous barcodes and trimming of adapter contamination were performed using Stacks version 2.1 (Catchen et al., 2013; Catchen et al., 2011).

Genome Analysis Toolkit (GATK) version 4.2.6.1 (Van der Auwera et al., 2013) was used to map paired-end and singleton ddRAD seq reads generated from pseudo-F1 sporophytes onto the *S. latissima* v2 genome assembly (Denoeud et al., 2024; https://phaeoexplorer.sb-roscoff.fr/) for SNP calling. The variant call format (VCF) file produced by GATK was then filtered with bcftools version 1.19 (Danecek et al., 2021) to remove SNPs that were potentially derived from multicopy genes, loci or samples with >20% missing data and loci with global minor allele frequency (MAF) of less than 0.01. Minimum allelic depth of 2 and coverage of 3 were required to keep a variant.

### Genetic diversity and population structure analyses

Prior to diversity and population structure analysis, SSR loci were tested for stuttering, large allele dropout and null alleles using the software MicroChecker v2.2.3 (Van Oosterhout et al., 2004). The frequency of null alleles per locus and per population was estimated according to the EM algorithm (Dempster et al., 1977) using FreeNA software (Chapuis & Estoup, 2007).

Genetic diversity was estimated per population, and expected heterozygosity (*H*e) and allelic richness (*Â)* were computed using the Hierfstat R package version 0.5-11 (Goudet & Jombart, 2024). The number of private allele (*PÂ*) was computed using the poppr R package version 2.9.8 (Kamvar et al., 2014) and standardized for the smallest sample sizes in terms of individuals within sites (N = 8), using 1000 randomizations.

Deviation from random mating was estimated by the inbreeding coefficient (*F*_IS_) for each population. *F*_IS_ was computed using the diveRsity R package (Keenan et al., 2013) version 1.9.90 and alleles were randomized 10,000 times among individuals within each sample to test for departure from random mating (*F*_IS_ significantly different from 0). The threshold for significance was set at p-value < 0.01.

To estimate the level of genetic differentiation between localities, pairwise *F*_ST_ values were computed in the Hierfstat R package version 0.5-11 (Goudet & Jombart, 2024). The significance of the comparisons was estimated by performing 10,000 bootstraps over loci (boot.ppfst function), with the comparisons judged significant if the bootstrap-generated confidence intervals did not overlap zero.

Genetic structure was inferred using Structure 2.3.4 (Pritchard et al., 2000) with admixture and a correlated allele frequency model, without any prior population assignments. A range of assumed populations (K, set sequentially from 1 to 12) was run 20 times using a burn-in of 5 x 10^5^ iterations and a run length of 1 x 10^6^ iterations. The number of clusters was estimated using the DeltaK criterion of Evanno et al. (2005). The Pophelper R package version 2.3.1 (Francis, 2017) was used to summarize assignment results across independent runs and to graphically visualize the results. CLUMPP version 1.1.2 (Jakobsson & Rosenberg, 2007) was used to align clustering outputs from multiple runs of STRUCTURE. The LargeKGreedy algorithm (M=3) was chosen with GREEDY OPTION=2, to test randomly 20 input orders of runs (REPEATS=20).

Structure analyses were completed with a discriminant analysis of principal components (DAPC) implemented in the adegenet R package version 2.1.11 (Jombart, 2008; Jombart et al., 2010). This analysis clusters individuals by refining genetic differentiation between populations while minimizing within-population differences.

### Heritability, GWAS, GS and putative candidate gene exploration

Marker-based heritability was estimated for each trait with the GCTA software (Yang et al., 2011) using the GREML method (Yang et al., 2010) to estimate the proportion of variance in a phenotype explained by all SNPs. GWAS for additive effects was performed for each trait with the BLINK (Bayesian-information and Linkage-disequilibrium Iteratively Nested Keyway) method implemented in the R GAPIT package version 3 (Lipka et al., 2012). The method links genotype to phenotype while minimizing false positives and increasing statistical power. It relies on two models: the first tests markers while including a subset of markers as cofactors, and the second iteratively selects the optimal set of markers (Huang et al., 2019). A minor allele frequency threshold of 1% was used and the first 3 PCA principal components were fitted as covariates to reduce the false positives due to population structure. Bonferroni threshold (0.05/number of used markers) was used to retain significant associations. Gene exploration was performed based on the genome annotation, considering the peak of significant associations or an interval of ± 100 kbp around the peaks.

GS ability was explored using the R BWGS package (Charmet et al., 2020). Prediction accuracy was estimated for each trait through a 5-fold cross-validation with 30 independent replicates. Eight prediction methods were tested: linear ridge-based model (GBLUP), penalized regression (LASSO and Elastic Net), Bayesian models (Bayes A, Bayes B, Bayes C and Bayesian LASSO) and nonlinear kernel model (RKHS).

## Results

### SSR genotyping and filtering, and efficacy of the 21 SSR loci to assign genotypes to their populations

A total of 197 SSR alleles were identified across the 13 sampled sites with 309 parental individuals and 537 pseudo-F1 individuals resulting in 846 unique multilocus genotypes (Supplementary Table 4). The mean number of alleles per locus was 8.14±2.16 (ranging from 2 to 16). There were 7.23% missing data across the final data set. MicroChecker analysis found no evidence of stuttering error or large dropout but did suggest the presence of null alleles in 13 of the 21 loci. For 5 loci (Sacl-11, Sacl-21, Sacl-33, Sacl-54, Sacl-60), null alleles were present in no more than two populations. For the remaining eight loci (Sacl-65, SLN32, SLN35, SLN36, SLN54, SLN319, SLN320, SLN510), null alleles were present in three to seven populations, but the average frequency of null alleles remained low i.e. <0.20 (Dakin & Avise, 2004). The average null allele frequency per locus ranged from 0.009±0.056 for Sacl-41 to 0.154±0.089 for SLN34 (see Supplementary Table 5).

The results showed that, for the chosen assignation threshold of score of 0.05, 99.4% (i.e: 307 of the 309 parental individuals, Supplementary Table 3, Figure 2) were correctly self-assigned to their population of origin. On this basis, these 21 SSR loci were used to assign pseudo-F1 to their population of origin with a 92.0% success rate (i.e: 494 among the 537 individuals genotyped, Supplementary Table 2, Figure 2). A total of 233 of the 494 pseudo-F1 that were successfully assigned were phenotyped and subsequently used for ddRAD-seq genotyping (Supplementary Table 6A and 6B, Figure 2).

### ddRAD-seq genotyping and single nucleotide polymorphisms (SNP) marker filtering

After demultiplexing, trimming and quality filtering the obtained reads, a total of 3.504 billion reads (81.9%) were retained, corresponding to an average of 14.9 million reads per sample and a mean coverage of 16x per sample. Based on this data, a total of 2,369,348 variants were called from 233 samples. After missing data, minor allele frequency and minimum allelic depth filtering, the final vcf file contained 202 samples with 148,065 biallelic variants for subsequent analyses (Figure 2).

### Genetic diversity and population structure

After filtering, the SSR dataset, comprising 21 loci with 164 alleles, and the SNP dataset, comprising 71,619 loci with 148,065 alleles across 202 pseudo-F1 individuals and 12 geographical sites, were used to conduct genetic diversity and population structure analyses (Table 1). The LOC site was excluded due to an insufficient sample size (n=1).

**Table 1.**
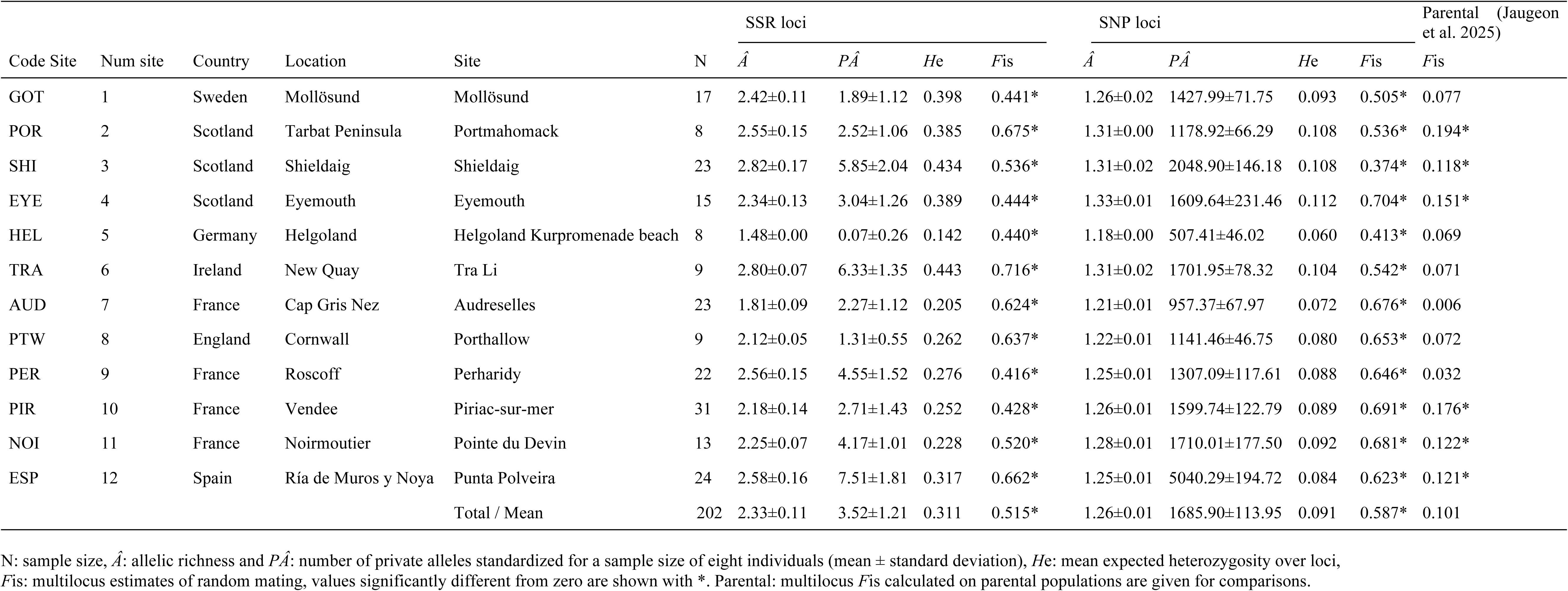
Genetic diversity of pseudo-F1 populations estimated with SSR loci and SNP loci.

| Code Site | Num site | Country | Location | Site | N | SSR loci |  |  |  | SNP loci |  |  |  | Parental (Jaugeon et al. 2025) |
| --- | --- | --- | --- | --- | --- | --- | --- | --- | --- | --- | --- | --- | --- | --- |
| | | | | | | $\hat{A}$ | $P\hat{A}$ | $H_e$ | $F_{is}$ | $\hat{A}$ | $P\hat{A}$ | $H_e$ | $F_{is}$ | |
| GOT | 1 | Sweden | Mollösund | Mollösund | 17 | 2.42±0.11 | 1.89±1.12 | 0.398 | 0.441* | 1.26±0.02 | 1427.99±71.75 | 0.093 | 0.505* | 0.077 |
| POR | 2 | Scotland | Tarbat Peninsula | Portmahomack | 8 | 2.55±0.15 | 2.52±1.06 | 0.385 | 0.675* | 1.31±0.00 | 1178.92±66.29 | 0.108 | 0.536* | 0.194* |
| SHI | 3 | Scotland | Shieldaig | Shieldaig | 23 | 2.82±0.17 | 5.85±2.04 | 0.434 | 0.536* | 1.31±0.02 | 2048.90±146.18 | 0.108 | 0.374* | 0.118* |
| EYE | 4 | Scotland | Eyemouth | Eyemouth | 15 | 2.34±0.13 | 3.04±1.26 | 0.389 | 0.444* | 1.33±0.01 | 1609.64±231.46 | 0.112 | 0.704* | 0.151* |
| HEL | 5 | Germany | Helgoland | Helgoland Kurpromenade beach | 8 | 1.48±0.00 | 0.07±0.26 | 0.142 | 0.440* | 1.18±0.00 | 507.41±46.02 | 0.060 | 0.413* | 0.069 |
| TRA | 6 | Ireland | New Quay | Tra Li | 9 | 2.80±0.07 | 6.33±1.35 | 0.443 | 0.716* | 1.31±0.02 | 1701.95±78.32 | 0.104 | 0.542* | 0.071 |
| AUD | 7 | France | Cap Gris Nez | Audreselles | 23 | 1.81±0.09 | 2.27±1.12 | 0.205 | 0.624* | 1.21±0.01 | 957.37±67.97 | 0.072 | 0.676* | 0.006 |
| PTW | 8 | England | Cornwall | Porthallow | 9 | 2.12±0.05 | 1.31±0.55 | 0.262 | 0.637* | 1.22±0.01 | 1141.46±46.75 | 0.080 | 0.653* | 0.072 |
| PER | 9 | France | Roscoff | Perharidy | 22 | 2.56±0.15 | 4.55±1.52 | 0.276 | 0.416* | 1.25±0.01 | 1307.09±117.61 | 0.088 | 0.646* | 0.032 |
| PIR | 10 | France | Vendee | Piriac-sur-mer | 31 | 2.18±0.14 | 2.71±1.43 | 0.252 | 0.428* | 1.26±0.01 | 1599.74±122.79 | 0.089 | 0.691* | 0.176* |
| NOI | 11 | France | Noirmoutier | Pointe du Devin | 13 | 2.25±0.07 | 4.17±1.01 | 0.228 | 0.520* | 1.28±0.01 | 1710.01±177.50 | 0.092 | 0.681* | 0.122* |
| ESP | 12 | Spain | Ría de Muros y Noya | Punta Polveira | 24 | 2.58±0.16 | 7.51±1.81 | 0.317 | 0.662* | 1.25±0.01 | 5040.29±194.72 | 0.084 | 0.623* | 0.121* |
| Total / Mean |  |  |  |  | 202 | 2.33±0.11 | 3.52±1.21 | 0.311 | 0.515* | 1.26±0.01 | 1685.90±113.95 | 0.091 | 0.587* | 0.101 |

N: sample size, $\hat{A}$ : allelic richness and $P\hat{A}$ : number of private alleles standardized for a sample size of eight individuals (mean ± standard deviation), $H_e$ : mean expected heterozygosity over loci, $F_{is}$ : multilocus estimates of random mating, values significantly different from zero are shown with \*. Parental: multilocus $F_{is}$ calculated on parental populations are given for comparisons.

A comparison of population genetic diversity for SSR loci and SNP loci is provided in Table 1. Results were very similar between the two types of markers except that the number of alleles and *H*e were higher for SSR loci compared to SNP loci, as expected, since SSR loci are multiallelic while SNP loci were bi-allelic.

Genotypic analysis using Structure (Figure 3, (a) for SSR loci and (b) for SNP loci) and DeltaK (Supplementary Figure 3, (a) for SSR loci and (b) for SNP loci) revealed two main clusters depending on the latitude: (1) northern Europe and (2) southern Europe, as expected (Jaugeon et al., 2025). The northern cluster contained sites from northern British Islands and Sweden, while the southern cluster grouped sites from Helgoland, the English Channel, Brittany and Spain (Figure 3). For SNPs, a substructure was observed within the southern clade (for K=4, DeltaK: Supplementary Figure 3; and structure graph: Figure 3) separating the North Sea populations from Spanish populations and a third group comprising populations from the English Channel and southern Brittany. The pattern observed with the DAPC analysis confirmed that observed with Structure. While the separation between the northern and southern clusters was found for SSRs (Supplementary Figure 4a, Supplementary Figure 5a), with SNPs (Supplementary Figure 4b), the populations of the northern cluster remained grouped together and the populations of the southern cluster differentiated into three groups according to the structure described above.

**Figure 3.**
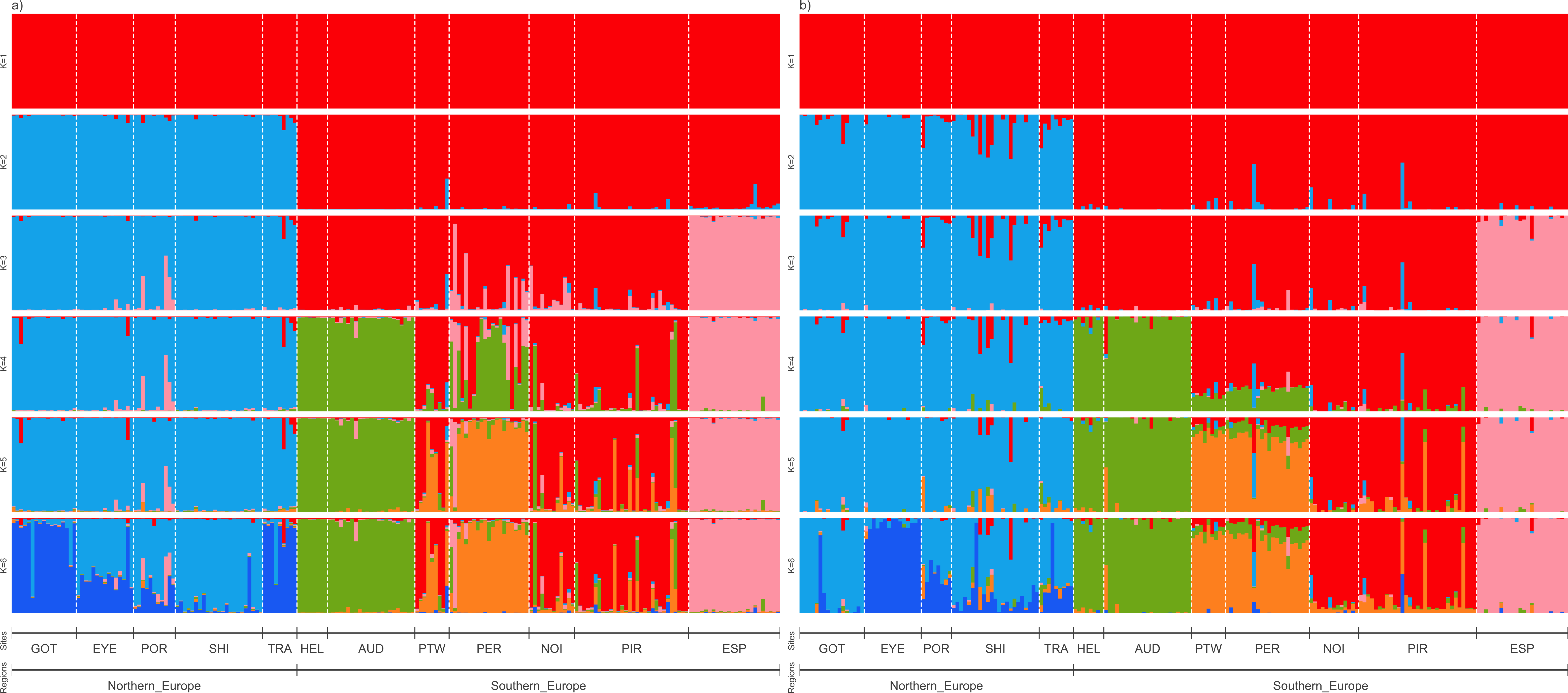
Genetic structure inferred by STRUCTURE from multi-locus SSR genotypes (a) and SNP genotypes (b). Hierarchical structure plots assuming K=1 (top) to K=6 (bottom) genotypic clusters. See Supplementary Table 1 for explanations of the location codes.

Pairwise *F*_ST_ values were significant between all sites and the pattern was very similar between SSR loci and SNP loci: values varied from 0.098 to 0.535 for SSR, and from 0.045 to 0.412 for SNP (Supplementary Figure 6). Mean *F*_ST_ values were also significant between northern and southern clusters with 0.264 for SSR loci and 0.207 for SNP loci.

Genetic diversity *(H*e) at SSR loci was higher for the sites of the northern cluster (from 0.385 in POR to 0.443 in TRA, Table 1) than those of southern cluster (from 0.142 in HEL to 0.276 in PER (Table 1), except in the extreme south where the *H*e increased to 0.317 in Spain (site ESP, Table 1). The same was true for SNP loci with *H*e values greater in the northern cluster (from 0.093 in sites GOT to 0.112 in EYE, Table 1) than in the southern cluster (from 0.06 in HEL to 0.092 in NOI, Table 1). For the northern cluster the *H*e values varied from 0.116 (SNP loci) to 0.471 (SSR loci) and for southern cluster from 0.100 (SNP loci) to 0.309 (SSR loci).

*F*_IS_ values were all highly significant compared to the parental population (Table 1), whatever the site and the type of genetic marker, as expected for pseudo-F1 individuals obtained by selfing (i.e.: expected *F*_IS_ value of 0.5, Table 1). For SSR loci, values varied from 0.416 (in PER) to 0.716 (in TRA) and from 0.374 (in SHI) to 0.704 (in EYE) for SNP loci (Table 1). Mean *F*_IS_ values over sites were 0.515 for SSR loci, very close to the value of 0.587 observed for the SNP loci. *F*_IS_ values were also all highly significant for both clusters and values varied from 0.585 (SSR loci) to 0.644 (SNP loci) for the northern cluster, and from 0.651 (SSR loci) to 0.674 (SNP loci) for the southern cluster.

### Morphological and metabolic traits of the pseudo-F1 population

The number of pseudo-F1 individuals measured for each morphological and metabolic trait per (assigned) site of origin is provided in Supplementary Table 6A and 6B, respectively. A total of 233 individuals were measured for each morphological trait whereas 131 individuals were analyzed for all metabolic traits, except for rhamnose concentration, which was measured in 113 individuals (Supplementary Table 7). Sample sizes varied considerably among populations, ranging from 2 to 34 individuals (Supplementary Table 6A and 6B). Only geographical sites represented by at least five individuals (i.e., five biological replicates) were retained for the statistical analyses i.e., 12 sites for morphological traits and 9 sites for all metabolic traits, except for rhamnose, for which only 8 sites were used (Supplementary Table 6A and 6B).

Boxplots of trait means for pseudo-F1 individuals at each assigned site are shown in Figure 4a for morphological traits and Figure 4b for metabolic traits. Generalized linear model (GLM) analyses revealed highly significant differences among sites for all morphological traits (Table 2A) and significant differences for all metabolic traits except rhamnose concentration (Table 2B). Overall, *p*-values were smaller for metabolic traits compared to morphological ones, except for amino-acid concentration showing highly significant differences among populations (Figure 4a and 4b for morphological and metabolic traits, respectively).

**Figure 4.**
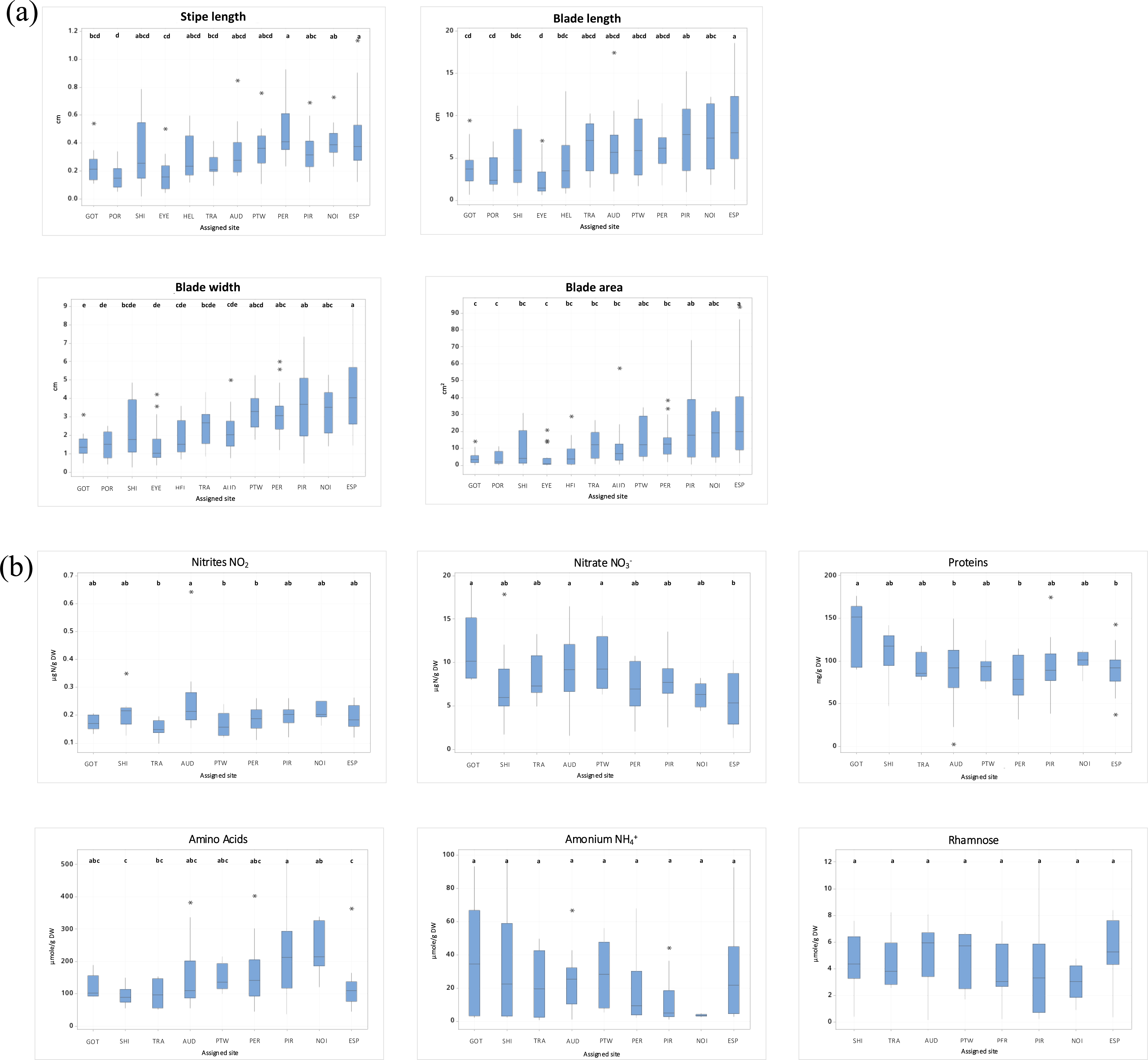
Box plots of the pseudo-F1 sporophytes assigned to each of the 12 geographical sites. Tukey pairwise comparisons of means was used for both morphological (a) and metabolic (b) traits. When letters are identical, the means are not significantly different.

**Table 2.** GLM analyses of the four morphological traits (2A) and the six metabolic traits (2B).

| Morphological traits | degree of freedom | Sum of square adjusted | Mean square adjusted | F-values | P-values |
| --- | --- | --- | --- | --- | --- |
| Stipe lenght (cm) | 11 | 1.91 | 0.17 | 6.5 | p<0.0001 |
| Blade length (cm) | 11 | 760.9 | 69.17 | 5.56 | p<0.0001 |
| Blade width (cm) | 11 | 200.8 | 18.26 | 9.65 | p<0.0001 |
| Blade surface aera (cm<sup>2</sup>) | 11 | 16351 | 1486.5 | 6.83 | p<0.0001 |

2B
| Metabolical traits | degree of freedom | Sum of square adjusted | Mean square adjusted | F-values | P-values |
| --- | --- | --- | --- | --- | --- |
| nitrate (NO<sub>2</sub>) (µgN/gDW) | 8 | 0.075 | 9 10<sup>-3</sup> | 2.92 | 0,005 |
| nitites (NO<sub>3</sub>) (µgN/gDW) | 8 | 258.4 | 32.3 | 3.16 | 0,003 |
| proteins (mg/gDW) | 8 | 14634 | 1829.3 | 2.46 | 0,017 |
| Amino Acids (µmole/gDW) | 8 | 269407 | 33676 | 4.2 | p<0.0001 |
| Amonium (NH<sub>4</sub>) (µmole/gDW) | 8 | 9078 | 1134.8 | 2.45 | 0,018 |
| Rhamnose (µmole/gDW) | 7 | 57.06 | 8.152 | 1.42 | 0.208 |

Overall, morphological traits exhibited a clear latitudinal pattern, with significantly higher values in southern populations than in northern ones (Figure 4a). For all four traits, mean values were significantly greater in the southernmost sites than in the northernmost sites, while intermediate values were observed at sites located close to the border between the two genetic clusters (HEL, TRA, AUD, PTW; Figure 4a). Higher variability was observed for SHI and ESP within the southern cluster, and for EYE within the northern cluster (Figure 4a). A Kruskal-Wallis test confirmed this latitudinal difference since each of the four morphological traits was always significantly higher in the southern than in the northern cluster (Supplementary Table 8A). In contrast, metabolic traits showed weaker spatial structuring and did not display a consistent latitudinal trend (Figure 4b). No significant differences among sites were detected for NH₄⁺ or rhamnose concentrations (Figure 4b). However, NO₃⁻ concentrations were significantly higher in GOT, AUD, and PTW compared with ESP. Protein concentrations were significantly higher in GOT than in ESP, PER, and AUD. NO₂ concentrations were higher in AUD than in TRA, PTW, and PER, while Amino Acid concentrations were significantly greater in PIR and NOI compared with SHI and ESP (Figure 4b). Kruskal-Wallis test did not show significant differences between clusters for most metabolic traits excepted for the amino acid concentration that was significantly higher in the southern than in the northern cluster (Supplementary Table 8B).

Phenotypic correlations were strong among morphological traits, as expected, reflecting coordinated variation in growth-related characteristics. In contrast, correlations between morphological and metabolic traits, as well as among metabolic traits, were generally weak or non-significant, suggesting relative independence between these trait groups. The main exceptions were significant associations between NO₃⁻ and NH₄⁺, and between NH₄⁺ and rhamnose (Supplementary Figure 7).

### QTL identifications and gene analysis

GWAS analysis with BLINK resulted in 26 significant associations at Bonferroni threshold of 3.38e^-7^ (alpha = 0.05), from five traits over the 10 traits evaluated (three morphological traits Table 3A and two metabolic traits, Table 3B; Figure 5; Supplementary Figure 8a-c and Supplementary Figure 8d-e respectively). Two significant associations were observed for blade length, three for blade width, 11 for blade area, three for NO_2_ and seven for NH_4_^+^. One association on chromosome 17 (SNP SL_17_8569693) was significant for both blade length and blade area.

**Figure 5.**
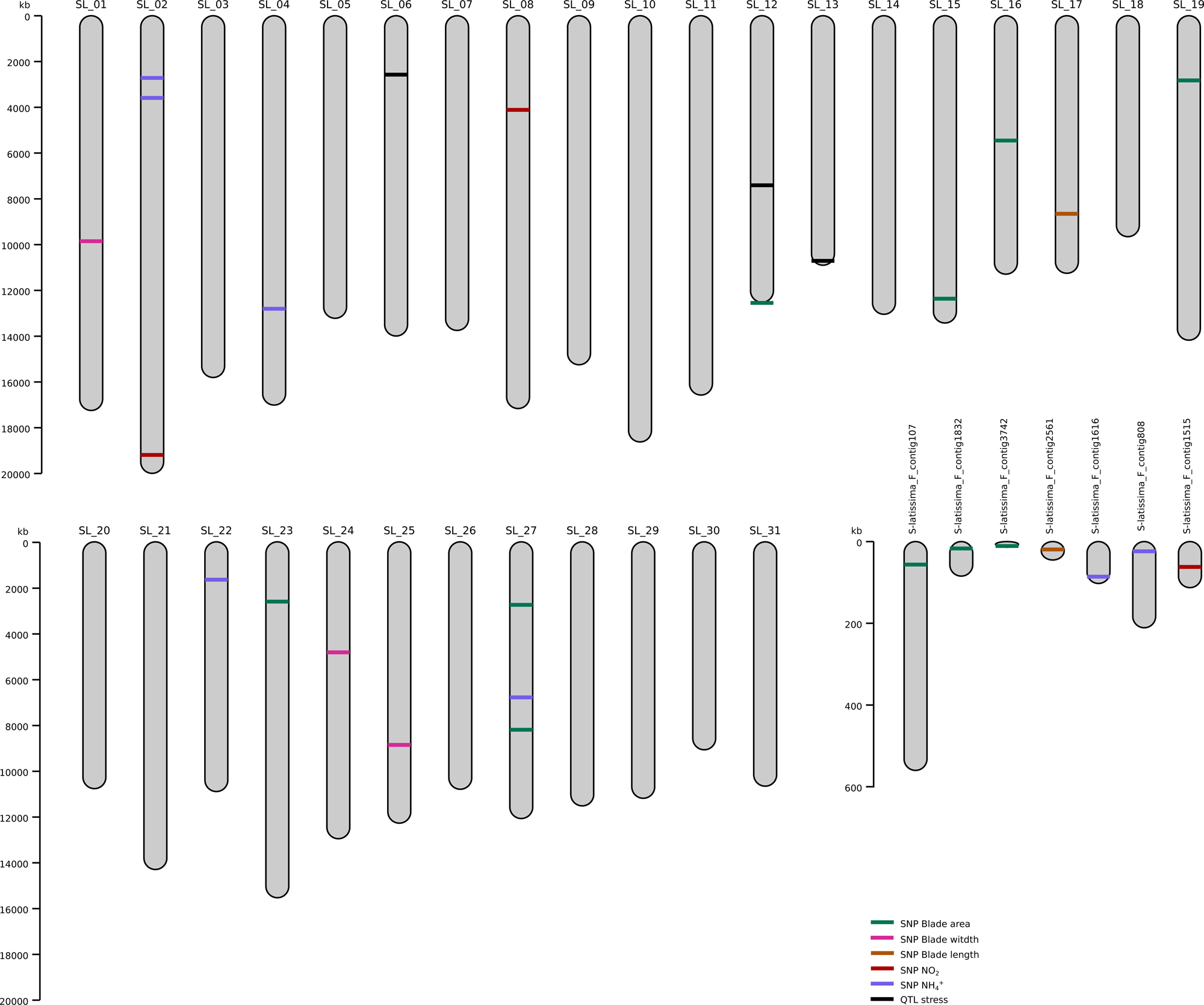
*S. latissima* genetic map showing the location of the QTLs associated to morphological and metabolic traits as well as temperature stress (Nehr et al., 2025).

**Table 3.** GWAS significant associations with morphological (A) and metabolic traits (B).

| Trait | SNP | Chromosome or contig | Position | *p*-value | MAF | Effect | PVE (%) |
| --- | --- | --- | --- | --- | --- | --- | --- |
| Blade length | S-latissima_F_contig2561_14541 | S-latissima_F_contig2561 | 14541 | 1.87E-07 | 0.054 | 3.522 | 27.61 |
| Blade length | SL_17_8569693 | SL_17 | 8569693 | 1.43E-09 | 0.172 | 1.996 | 25 |
| Blade width | SL_01_9768174 | SL_01 | 9768174 | 4.75E-08 | 0.012 | -1.66 | 52.44 |
| Blade width | SL_24_4743110 | SL_24 | 4743110 | 1.26E-08 | 0.079 | -0.643 | 2.005 |
| Blade width | SL_25_8786509 | SL_25 | 8786509 | 2.14E-19 | 0.052 | -1.345 | 23.71 |
| Blade area | S-latissima_F_contig107_51635 | S-latissima_F_contig107 | 51635 | 2.39E-17 | 0.039 | 11.17 | 4.956 |
| Blade area | S-latissima_F_contig1832_12152 | S-latissima_F_contig1832 | 12152 | 1.11E-08 | 0.167 | 3.957 | 0.651 |
| Blade area | S-latissima_F_contig3742_6231 | S-latissima_F_contig3742 | 6231 | 4.82E-08 | 0.057 | 5.926 | 7.354 |
| Blade area | SL_12_12462931 | SL_12 | 12462931 | 1.16E-10 | 0.293 | 3.787 | 0.943 |
| Blade area | SL_15_12279389 | SL_15 | 12279389 | 3.08E-13 | 0.017 | 13.022 | 7.091 |
| Blade area | SL_16_5368748 | SL_16 | 5368748 | 5.96E-13 | 0.03 | 10.741 | 7.385 |
| Blade area | SL_17_8569693 | SL_17 | 8569693 | 1.91E-11 | 0.172 | 5.012 | 1.866 |
| Blade area | SL_19_2742247 | SL_19 | 2742247 | 1.88E-11 | 0.101 | 7.662 | 2.624 |
| Blade area | SL_23_2526597 | SL_23 | 2526597 | 6.67E-23 | 0.015 | 22.904 | 16.747 |
| Blade area | SL_27_2667616 | SL_27 | 2667616 | 6.21E-34 | 0.01 | -47.287 | 45.218 |
| Blade area | SL_27_8124811 | SL_27 | 8124811 | 2.20E-09 | 0.059 | 6.206 | 2.8 |

B)
| Trait | SNP | Chromosome or contig | Position | *p*-value | MAF | Effect | PVE (%) |
| --- | --- | --- | --- | --- | --- | --- | --- |
| NO<sub>2</sub> | S-latissima_F_contig1515_57359 | S-latissima_F_contig1515 | 57359 | 1.45E-10 | 0.137 | 0.04 | 20.706 |
| NO<sub>2</sub> | SL_02_19104822 | SL_02 | 19104822 | 1.94E-09 | 0.057 | 0.046 | 39.87 |
| NO<sub>2</sub> | SL_08_4030272 | SL_08 | 4030272 | 6.98E-09 | 0.17 | -0.026 | 16.66 |
| NH<sub>4</sub><sup>+</sup> | S-latissima_F_contig1616_81148 | S-latissima_F_contig1616 | 81148 | 2.83E-09 | 0.078 | 14.317 | 7.466 |
| NH<sub>4</sub><sup>+</sup> | S-latissima_F_contig808_19157 | S-latissima_F_contig808 | 19157 | 9.64E-14 | 0.119 | 15.294 | 8.508 |
| NH<sub>4</sub><sup>+</sup> | SL_02_2634186 | SL_02 | 2634186 | 7.92E-11 | 0.087 | 16.257 | 6.112 |
| NH<sub>4</sub><sup>+</sup> | SL_02_3507683 | SL_02 | 3507683 | 5.39E-15 | 0.037 | 28.524 | 35.067 |
| NH<sub>4</sub><sup>+</sup> | SL_04_12718968 | SL_04 | 12718968 | 1.03E-09 | 0.069 | 17.676 | 9.41 |
| NH<sub>4</sub><sup>+</sup> | SL_22_1567909 | SL_22 | 1567909 | 2.08E-08 | 0.22 | 6.008 | 1.476 |
| NH<sub>4</sub><sup>+</sup> | SL_27_6710473 | SL_27 | 6710473 | 1.31E-16 | 0.041 | 25.602 | 28.403 |

MAF: minor allele frequency; Effect: allelic effect of the minor allele at the genotype; PVE: percentage of variance explained by the genotype.

We found seven genes under the association peaks based on the genome annotation. The seven genes are putatively involved in ATPase activity, GTP binding, regulation of response to stimulus, peptidyl-lysine acetylation, axoneme assembly, ATP-binding and glycosylase activity (Table 4). To take into account linkage disequilibrium, the search region was enlarged to ± 100 Kbp around the association peaks. This identified 129 unique genes for the 5 traits with significant associations (Supplementary Table 9).

**Table 4.**
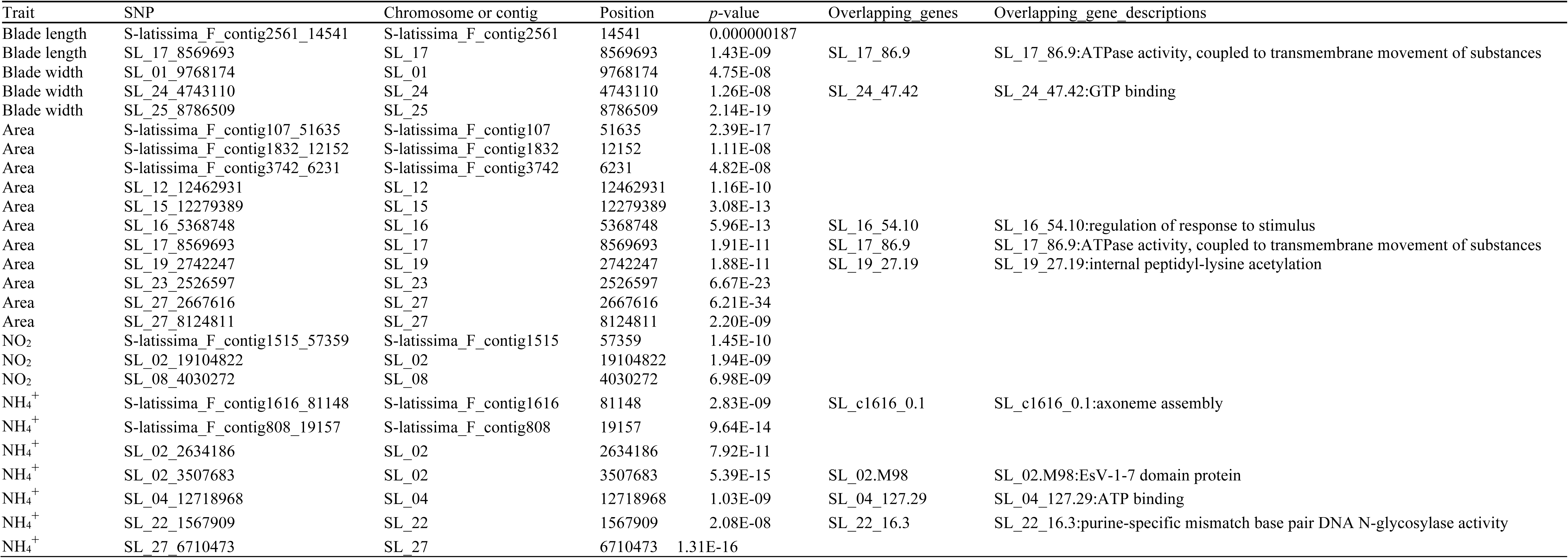
Annotated genes under GWAS association peaks with the 5 traits.

Several significant loci explained a substantial proportion of the phenotypic variance (PVE). For blade length, locus SL_01_9768174 accounted for 52.44% of the variance, while three additional loci associated with the same trait showed PVEs ranging from 23.71% to 27.61%. Blade area was also strongly associated with locus SL_27_2667616, which explained 45.21% of the variance. For metabolic traits, NO₂ exhibited two significant peaks with PVEs of 29.87% and 20.70%, whereas NH₄^+^ showed two peaks explaining 35.06% and 28.40% of the variance.

With respect to morphological traits, most detected loci had positive effects. An exception was blade width, for which all three associated loci showed negative effects, and one of the eleven loci identified for blade area had a pronounced negative effect. Similarly, among metabolic traits, only a single SNP associated with NO₂ showed a negative effect.

### Heritability and predictive ability

Marker-based heritability values were high for all morphological traits, while metabolic traits showed moderate to low heritability, except for NH₄⁺. Cross-validation prediction’s accuracy was moderate to high for most traits but negligible for protein content. Overall, morphological traits exhibited higher predictive ability than metabolic traits (Table 5, Supplementary Figure 9).

**Table 5.** Marker-based heritability values and predictive abilities.

| Trait | Heritability $h^2$ (SE) | Best prediction method | Predictive ability |
| --- | --- | --- | --- |
| Stipe | 0.75 (0.26) | GBLUP | 0.45 |
| Total length | 0.89 (0.29) | EN | 0.49 |
| Blade length | 0.89 (0.29) | EN | 0.49 |
| Blade width | 0.99 (0.30) | RKHS | 0.59 |
| Blade area | 0.99 (0.31) | RKHS | 0.54 |
| NO<sub>2</sub> | 0.13 (0.24) | BC | 0.23 |
| NO<sub>3</sub><sup>-</sup> | 0.14 (0.18) | RKHS | 0.26 |
| Proteins | 0.00 (0.17) | BC | -0.007 |
| Amino acids | 0.40 (0.30) | BA | 0.41 |
| NH<sub>4</sub><sup>+</sup> | 0.99 (0.52) | BL | 0.21 |
| Rhamnose | 0.20 (0.28) | EN | 0.35 |

Heritability values were given with their standard errors in parentheses.

## Discussion

### Experimental design considerations

Our use of self-fertilized pseudo-F1 progeny from 13 wild populations spanning northern and southern European genetic clusters offers both advantages and limitations for GWAS. Selfing creates controlled genetic backgrounds that reduce environmental variance and increase power to detect genetic effects, analogous to the use of recombinant inbred lines or doubled haploids in crop genetics. The high assignment success of pseudo-F1 individuals to their parental populations confirms maintenance of population-level genetic structure and validates the use of population-of-origin as a covariate in GWAS models. However, achieving our final sample size of 202 individuals required navigating substantial experimental challenges, as illustrated in Figure 2. The workflow from parental populations through gametophyte culture, fertilization, and sporophyte development involved significant attrition due to gametophyte contamination, failed fertilization events, early sporophyte mortality, and losses during phenotyping. These challenges are inherent to kelp experimental systems, where the complex biphasic life cycle, sensitivity to culture conditions, and requirement for precise environmental control during gametogenesis impose practical constraints on population sizes (Bråtelund et al., 2026). Such limitations make outcrossed designs, which would require controlled crosses between specific gametophyte pairs, even more logistically demanding, partly justifying our selfing approach despite its drawbacks.

### Significant genetic differentiation among populations of origin and strong divergence between northern and southern European clusters

Although SSR markers provided slightly higher resolution than SNP markers, the two marker types yielded congruent results and consistently revealed two major genetic clusters of *S. latissima* across Europe, corresponding to northern and southern regions. This north-south genetic structure has been repeatedly detected using a range of mitochondrial and nuclear molecular markers (Diehl et al., 2023; Guzinski et al., 2016; Guzinski et al., 2020; Nehr et al., 2025; Neiva et al., 2018). Notably, a similar clustering pattern was previously described by Neiva et al. (2018) using SSR markers alone. Comparable concordance between SSRs and SNPs in the estimation of genetic diversity and population structure has been reported in multiple species, including kelps (Guzinski et al., 2020; Sunde et al., 2020). The pattern of genetic diversity and population structure was also congruent between the two types of markers. Higher within-population genetic diversity and weaker population structure in the northern populations compared with southern populations support the hypothesis of contrasting postglacial recolonization scenarios for the two clusters: recolonization from a northern refugium for the northern cluster, and recolonization from multiple refugia located near Brittany and Spain for the southern cluster (Jaugeon et al., 2025). The hypothesis of a glacial refugium in northern Europe with individuals persisting through the Last Glacial Maximum in Arctic or sub-Arctic regions, is relatively recent but is gaining increasing support for several cold-water species, including kelps (Bringloe et al., 2020; Fragkopoulou et al., 2025).

The pronounced genetic divergence between the two clusters, together with their distinct demographic histories, has likely shaped their evolutionary response to selection. Accordingly, we expected to identify loci associated with this genetic differentiation in the present study. Furthermore, within the southern cluster, the observed genetic structure separating North Sea populations from Spanish populations, with an intermediate group comprising populations from the English Channel and southern Brittany, suggests the potential for additional differentiation in morphological and metabolic traits among populations.

Consistent with these expectations, we observed significant differences in morphological traits between northern and southern genetic clusters. Pseudo-F1 individuals originating from the southern cluster were consistently larger than those from northern Europe, indicating divergence in growth-related traits that may reflect both historical and local adaptation.

### Significant difference between F1-pseudo populations for morphological traits related to latitude

Morphological plasticity is widespread in *S. latissima* and is strongly influenced by environmental conditions (Diehl et al., 2024). For instance, exposure to strong wave action and currents promotes the development of narrow-bladed sporophytes (Zhu et al., 2021). However, accumulating experimental evidence indicates that morphological variation in this species is not exclusively environmentally driven and that a substantial genetic component is involved. Marked morphological differences have been reported between populations from Brittany (Roscoff, France) and the Arctic (Svalbard, Norway), with Arctic sporophytes being longer and narrower, whereas Brittany individuals are wider and shorter (Monteiro et al., 2021). Moreover, Arctic sporophytes exhibited faster growth rates than those from Brittany under laboratory conditions, and population-specific morphologies were maintained in sporophytes derived from cultured gametophytes, supporting a genetic basis for these traits (Monteiro et al., 2021). Similarly, studies conducted along the coasts of Devon (southern UK) and Argyll (Scotland) showed that an increase of approximately 6° in latitude was associated with higher frond growth rates and a developmental delay of more than a month at the northern site (Parke, 1948). These differences were interpreted as being at least partly genetically determined. Comparable patterns have also been observed between glacial and oceanic populations in Alaska, where reciprocal transplant experiments in Kachemak Bay demonstrated that thalli largely maintained the seasonal growth patterns characteristic of their native habitats (Spurkland & Iken, 2012). By combining genome-scale data with field-measured growth rates, Starko et al. (2026) have evidenced a genomic basis for growth rate variation in a Western Australian population of *Ecklonia radiata*. Together, these results suggest that variation in morphology and growth in *S. latissima* reflects, at least in part, genetically based differentiation, likely representing adaptive responses to local environmental conditions. More recently, transcriptomic analyses have reinforced this conclusion by revealing a high number of differentially expressed genes (DEGs) between sporophytes from Brittany and the Arctic (Machado Monteiro et al., 2019).

Although Arctic populations were not included in the present study, the comparison between two genetically distinct European groups, northern and southern, provides further insight into the genetic basis of morphological differentiation. Our results clearly showed significant differences in all measured morphological traits between these two clusters. However, in contrast to previous studies (Monteiro et al., 2021; Parke, 1948), sporophytes from the northern group were smaller than those from the southern group. This pattern suggests that pseudo-F1 individuals from the southern cluster may exhibit higher growth rates at juvenile stages than those from the northern cluster. Importantly, substantial variability was also observed among populations within clusters, potentially reflecting fine-scale local adaptation linked to geographic location or latitude. Growth and reproductive strategies in kelps are known to depend on both environmental cues and endogenous regulatory mechanisms (Diehl et al., 2024), including circannual clocks synchronized by photoperiodic cycles (Lüning, 1993). The differences between our results and those of earlier studies may therefore reflect differences in life stage, as morphological traits in the present study were measured in juvenile sporophytes, whereas previous studies primarily focused on adult individuals.

An alternative, but not mutually exclusive, explanation involves genetic effects associated with selfing. All morphological traits were measured 90 days after self-fertilization of the parental sporophytes. We hypothesize that the larger size of pseudo-F1 individuals from the southern cluster may partly result from weaker inbreeding depression affecting sporophyte size or growth than in the northern cluster. Northern parental populations exhibited higher genetic diversity and larger effective population sizes, better masking and accumulating more deleterious mutations, which may render their self-fertilized progeny more susceptible to inbreeding depression. Consistent with this interpretation, significant inbreeding depression following selfing has been reported in *S. latissima* for the same morphological traits analyzed here (stipe length, blade length, and blade width) in a three-generation selection experiment using parents sampled from a wild Norwegian population (Bråtelund et al., 2026).

### Difference between F1-pseudo populations for metabolic traits

In contrast to clear morphological differences between northern and southern cluster pseudo-F1 populations, metabolic traits showed more complex patterns. While ANOVA results revealed significant population effects for some metabolic traits, these differences did not consistently align with the north-south geographic clustering observed for morphology, suggesting different genetic architecture and evolutionary drivers. One explanation is that metabolic traits related to nitrogen accumulation may reflect local environmental adaptation rather than broad latitudinal patterns. Nitrogen availability varies substantially across the European range of *S. latissima*, with differences in upwelling intensity, nutrient regimes, and seasonal patterns between Atlantic, North Sea, and Baltic populations (Gevaert et al., 2001; Nielsen et al., 2014). In the other side of the Atlantic in Canada, strong genetic differentiation between the populations of sugar kelp (*Laminaria longicruris*, now *S. latissima*) has been demonstrated in nitrate-poor (St. Margaret’s Bay) and nitrate-rich (Bay of Fundy) regions of Nova Scotia (Espinoza & Chapman, 1983). Maximum specific growth rate (μmax) and maximum uptake rate (Vmax) for nitrate were higher for St. Margaret’s Bay than for Fundy plants. Genetic variation for metabolic traits might therefore reflect a mosaic of local adaptations to site-specific nutrient conditions rather than simple geographic structure. Alternatively, metabolic traits may be more environmentally sensitive than morphological traits. Multi-environment trials across different nutrient regimes could help disentangle genetic effects from genotype-by-environment interactions.

The year-round protein concentration and amino acid content of maricultured *S. latissima* were determined in several previous studies in the Faroe Islands (Grandorf-Bak et al., 2019) and in Denmark (Marinho et al., 2015). These studies showed lower concentration in the upper latitude pointing on a potential genetic differentiation that may be correlated with growth strategies that are strongly dependent of the light availability and the summer temperature in the different regions that could also be related to local adaptations.

### GWAS and gene analysis

This study represents one of the first comprehensive GWAS analyses in *S. latissima* and contributes to the emerging genomic toolkit for kelp breeding. To date, GWAS and QTL mapping efforts in macroalgae remain limited compared to terrestrial crops, but recent advances in kelp and other seaweed species provide a valuable comparative context for interpreting our results. In the red alga *Gracilariopsis lemaneiformis*, GWAS of 174 haploid strains identified candidate genes associated with growth rate and branching, including genes involved in cell cycle regulation (cell division cycle protein 20-like), transcriptional control (DPB-like transcription factor), and auxin transport (Auxin transport protein BIG) (Feng et al., 2023). Similarly, QTL mapping in *Pyropia haitanensis* using SLAF-seq markers identified loci for morphological traits with individual QTL explaining 9.59-16.61% of phenotypic variance (Xu et al., 2015). In the kelp *S. japonica*, multiple QTL studies have mapped blade length and width using AFLP, SSR, and SNP markers, with collective QTL variance explained ranging from 36.39% for frond width to 42.36% for frond length (Liu et al., 2010; Wang et al., 2018).

Our study detected 26 significant associations across five traits (blade length, blade width, blade area, NO₂, and NH₄^+^) at stringent Bonferroni thresholds. The phenotypic variance explained by individual loci ranged from 0.65% to 52.44%, with several major-effect loci explaining >20% of trait variance. These PVE estimates exceed those typically reported in most previous algal GWAS studies and are comparable to major QTL detected in biparental mapping populations of crops. For instance, in rice recombinant inbred lines, major grain size QTLs explained 65.2-72.5% of variance (Kabange et al., 2023), while in maize doubled haploids, individual QTL for yield components under stress explained 4.4-19.4% (with one reaching 37.3%) (Cerrudo et al., 2018). The relatively high PVE values observed here likely reflect the pseudo-F1 population structure derived from self-fertilized sporophytes, which creates a controlled genetic background similar to biparental crosses while maintaining some natural diversity across the 12 source populations. As a result, effect sizes may be inflated compared with those obtained from more heterogeneous natural populations. A notable association peak on chromosome 17 (SNP SL_17_8569693) was shared between blade length and blade area, explaining 25% and 1.87% of the variance, respectively. These traits were strongly correlated in the pseudo-F1 populations. The co-localization of association signals across correlated traits is commonly observed in crop GWAS and may reflect either pleiotropy or close physical linkage between causal variants (Bretani et al., 2022; Zhao et al., 2019). Given that blade area is largely a function of blade length, one can assume that the same causal variant is involved in their expression. In contrast, blade width was associated with a different set of loci, suggesting partially independent genetic control of different dimensions of blade morphology. This pattern may indicate modular genetic architecture underlying blade development. Alternatively, given the strong phenotypic correlation between these traits, it could reflect differences in statistical power, with some shared loci remaining undetected. Although trade-offs in resource allocation during blade growth where selection for increased width comes at a cost to other growth parameters are plausible, the current data do not allow this hypothesis to be tested directly. Overall, these results highlight a genetic architecture combining major-effect loci and trait-specific associations, while also emphasizing the influence of population design on effect size estimates and the need for further validation in independent populations.

We identified seven genes directly under association peaks and 129 additional genes within ± 100 kbp windows. Genes at peaks included functions related to membrane transport, intracellular trafficking, cytoskeletal dynamics, and regulatory processes, with additional contributions from primary metabolism and protein turnover. This suggests that variation in blade morphology and nitrogen-related traits in *S. latissima* is primarily governed by mechanisms controlling cell expansion, intracellular organization, and resource allocation, rather than by single pathways of nutrient assimilation alone. The association of blade dimensions with an ATPase gene involved in transmembrane transport is intriguing, as active transport processes are critical for nutrient uptake and turgor-driven cell expansion (Hurd et al., 2014). Variation in ATPase activity may modulate nutrient acquisition efficiency during rapid blade expansion. These interpretations remain however speculative without direct experimental validation through transcriptomic analyses and functional assays.

### Heritability estimates and genetic architecture

The marker-based heritability estimates for morphological traits in our study were exceptionally high, while metabolic traits showed a broader range, with proteins showing no heritability and NH₄^+^ showing h² = 0.99. However, these estimates must be interpreted with caution. The very large standard errors associated with our heritability estimates (ranging from 0.17 to 0.52) indicate substantial uncertainty in these values and reflect well-known limitations of marker-based heritability estimation in populations of moderate size (de los Campos et al., 2015; Kruijer et al., 2015; Speed et al., 2017). Marker-based heritabilities are prone to errors arising from several sources, including sampling variance due to limited population size, which can lead to both over- and underestimation of true heritabilities, population structure, which can inflate heritability estimates, linkage disequilibrium patterns specific to the study population, which may not generalize to other populations or breeding programs, and incomplete marker coverage, particularly for structural variants or regulatory regions not well-tagged by SNPs (Visscher et al., 2008; Wray et al., 2013). The pseudo-F1 design used in our study, while offering advantages for QTL detection, may further complicate heritability estimation due to the confounding effects of inbreeding and the limited number of recombination events compared to outbred populations (Sillanpää, 2011). Given these limitations, our heritability estimates should be viewed as preliminary indicators of the potential for genetic improvement rather than definitive quantifications of trait genetic control. They provide a practical starting point for assessing which traits are likely to respond to selection, but further studies with larger, independent breeding populations will be necessary to obtain robust, generalizable heritability estimates for *S. latissima*. A recent QTL study in the species reported significant inbreeding depression for stipe length, blade length, and blade width, and only stipe length had a similar range of heritability to our study, the other traits showing lower values (Bråtelund et al., 2026). The near-zero heritability for protein content and low heritability estimates for NO₂ and NO₃^-^ suggest that these metabolic traits are either more strongly influenced by environmental factors in our common garden setup, have limited standing genetic variation in the populations sampled, or that our marker panel does not adequately capture the genetic variation underlying these traits.

### Genomic selection: cross-validation results and future prospects

To complement the multi-locus GWAS approach and assess the collective predictive power of genome-wide markers, we performed GS analyses using multiple prediction models. It is important to emphasize that these genomic selection results represent cross-validation within the current pseudo-F1 population and should be interpreted as exploratory analyses that provide an initial indication of the potential for genomic prediction in *S. latissima*. Cross-validation assesses prediction accuracy within the training population tends to overestimate the accuracy achievable in independent breeding populations due to population-specific LD patterns, relatedness structure, and potential overfitting to the training data (Habier et al., 2007; Wientjes et al., 2013). Definitive evaluation of genomic selection for kelp breeding will require validation in independent forward-prediction scenarios, where models trained on one reference population or generation are used to predict phenotypes in subsequent breeding cycles or unrelated populations. Nevertheless, the cross-validation results offer several valuable insights. First, the fact that genomic selection models achieved moderate-to-high predictive abilities (0.45 - 0.59) for all morphological traits, including stipe length which showed no significant GWAS associations, supports the hypothesis that these traits have polygenic architectures with many small-effect loci collectively captured by genome-wide markers (Meuwissen et al., 2001). This “missing heritability” phenomenon, where traits exhibit high marker-based heritability but few individually significant GWAS hits, is common in both crop and animal genetics and motivates the use of genomic selection to harness the cumulative effects of undetected loci (Manolio et al., 2009). The variation in predictive ability across traits and prediction methods provides clues about genetic architecture. Traits with the highest predictive abilities tended to favor either kernel-based methods (RKHS, GBLUP) or regularized regression (Elastic Net), suggesting a combination of additive polygenic effects and potentially non-additive interactions (de los Campos et al., 2010). In contrast, traits with lower predictive abilities may have genetic architectures poorly captured by the current marker panel, stronger environmental components, or measurement error issues (Goddard et al., 2009). The comparison between GWAS and genomic selection highlights complementary strengths. GWAS excels at identifying specific large-effect loci suitable for marker-assisted selection (e.g., the blade width locus explaining 52.44% PVE and the blade area locus explaining 45.22% PVE), while genomic selection leverages all marker information to predict breeding values for selection decisions (Heffner et al., 2009). For traits with mixed architectures, integrating GWAS-detected QTL as fixed effects or weighted markers in genomic selection models can improve prediction accuracy, as demonstrated in poplar and wheat (Arruda et al., 2015; Guo et al., 2025). For kelp aquaculture, where generation times are currently 1-2 years and phenotyping at sea is logistically challenging and expensive, genomic selection could similarly accelerate breeding by enabling early selection and reducing the need for extensive field trials (Goecke et al., 2020). In summary, while our genomic selection results are preliminary and require validation in independent breeding populations, they demonstrate proof-of-concept that genome-wide markers can predict morphological trait performance in *S. latissima* with moderate accuracy. Combined with the major-effect loci identified by GWAS, these findings lay the groundwork for a hybrid breeding strategy that integrates marker-assisted selection for large-effect loci with genomic selection for polygenic background, maximizing genetic gain across economically important traits.

### Sample size, statistical power, and detection of genetic architecture

The GWAS analysis used 202 pseudo-F1 individuals genotyped at 148,000 SNP markers, representing a moderate sample size by crop GWAS standards but among the largest for any kelp genetic study to date. Sample size is a critical determinant of statistical power to detect associations, particularly for loci with small effect sizes (Spencer et al., 2009). In crops, large diversity panels with thousands of individuals can detect subtle associations (PVE <5%), whereas smaller studies are powered primarily to detect major-effect loci. The fact that we detected 26 significant associations with PVE values ranging from 0.65% to 52.44% suggests our sample size was adequate for discovering moderate-to-large effect loci but likely underpowered for small-effect variants. This is consistent with the “missing heritability” discussed earlier: traits with high marker-based heritability but few GWAS hits probably have polygenic architectures with many undetected small-effect loci. The success of genomic selection models in predicting these traits (predictive abilities 0.45-0.59) supports this interpretation, as genomic selection captures collective effects of all markers, including those below the significance threshold (Meuwissen et al., 2001). Interestingly, traits with the most significant associations (blade area: 11 loci; NH₄^+^: 7 loci) also showed very high heritability (h² = 0.99 for both, though with large standard errors), suggesting that substantial genetic variance for these traits is captured by detectable loci. This pattern resembles a mixed genetic architecture with major loci plus moderate-effect loci.

### Implications for kelp breeding: integrating GWAS and genomic selection

From a breeding perspective, the combination of GWAS loci and genomic selection results provides a practical foundation for genetic improvement in *S. latissima*. The major-effect loci for blade morphology (e.g., SL_01_9768174 for blade width with 52.44% PVE, SL_27_2667616 for blade area with 45.22% PVE) are particularly attractive targets for marker-assisted selection (MAS), as they explain large proportions of variance and should be reliably detected across breeding populations (Collard & Mackill, 2008). However, given the polygenic architecture evident for some traits and the moderate predictive abilities achieved by genomic selection models, a hybrid breeding strategy integrating both approaches is likely most effective. For traits controlled by large-effect loci (blade width, blade area), MAS can efficiently fix favorable alleles in breeding populations. For traits with high heritability but few GWAS hits (stipe length, blade length), genomic selection offers a path to capture cumulative effects of many small-effect loci (Meuwissen et al., 2001). Implementing GS will require training populations of 300-500 individuals and integration with high-throughput phenotyping workflows (Voss-Fels et al., 2019). Based on crop experiences, GS could achieve 15-25% genetic gains per cycle while reducing generation intervals (Cerrudo et al., 2018). An optimal hybrid strategy would use MAS to fix favorable alleles at major loci while simultaneously applying genomic selection genome-wide to improve polygenic backgrounds (Bernardo, 2014; Spindel et al., 2016).

### Challenges and future directions

Several challenges and opportunities emerge from this study. First, the use of self-fertilized pseudo-F1 progeny, while enabling rapid population generation, introduces inbreeding depression that may confound trait values and complicate interpretation of geographic patterns. Future studies should compare self-fertilized and outcrossed families to partition genetic effects from inbreeding effects and assess the magnitude of heterosis in kelp hybrids (Cohen et al., 2025). Second, functional annotation of kelp genomes remains incomplete, and many associated SNPs fall in unannotated regions or genes with unknown functions. Expanding functional genomic resources, including tissue-specific transcriptomes, proteomes, and metabolomes, will be essential for moving from statistical associations to biological understanding. Integration of multi-omics data can help prioritize candidate genes; for instance, combining GWAS with expression QTL (eQTL) mapping can identify variants that alter transcript abundance and link them to downstream phenotypic effects. Third, the genetic architecture of traits may vary between juvenile sporophytes (studied here at 90 days post-fertilization) and mature, commercially relevant life stages. Longitudinal phenotyping and age-specific GWAS would clarify temporal dynamics of trait expression. Fourth, our study focused on controlled common-garden conditions, but kelp aquaculture operates in diverse ocean environments. The results obtained in laboratory well controlled conditions might not necessarily reflect properly the performance of given individuals when grown for a longer period in the unstable environment they would face at sea. In line, a study using Arabidopsis plants grown either outside or in well controlled laboratory conditions showed a relatively poor correlation (r^2^ = 0.26) for phenotypic traits between both environments (Poorter et al., 2016). Understanding genotype-by-environment interactions (GxE) and identifying loci with stable effects across environments versus those conferring local adaptation will be critical for breeding broadly adapted or environment-specific cultivars (Des Marais et al., 2013). Multi-environment GWAS and genomic prediction models incorporating environmental covariates represent promising approaches for dissecting GxE (Jarquín et al., 2014).

Fifth, the large standard errors on heritability estimates and the preliminary nature of genomic selection cross-validation results underscore the need for larger, more diverse breeding populations to obtain robust genetic parameter estimates. Collaborative efforts across kelp research groups to pool data and establish multi-site breeding trials could accelerate progress and improve generalization of findings.

## Conclusion

This study demonstrates the power and potential of GWAS for dissecting the genetic basis of economically important traits in *S. latissima*. We identified 26 significant associations across morphological and metabolic traits, with several major-effect loci explaining substantial phenotypic variance (up to 52.44% for blade width). Genomic selection analyses, while preliminary and requiring validation in independent breeding populations, suggest that genome-wide markers can predict trait performance with moderate accuracy (0.45-0.59), particularly for polygenic traits lacking significant GWAS hits. Comparative analysis with other algal and crop GWAS studies reveals that our findings are consistent with emerging patterns in kelp genomics and provide a foundation for integrated breeding strategies combining marker-assisted selection and genomic selection. The marker-based heritability estimates, though subject to large standard errors and requiring cautious interpretation, provide a practical starting point for assessing the potential for genetic improvement. Further studies with larger, independent breeding populations and multi-environment trials will be essential to refine these estimates and validate the breeding utility of identified loci and genomic prediction models.

### Funding

This work benefited from the support of the French Government through the National Research Agency with regards to an investment expenditure program IDEALG (ANR-10-BTBR-04), the EU Horizon 2020 project GENIALG (Grant Agreement No 727892).

## Acknowledgments

We are grateful to the Roscoff Bioinformatic platform (ABiMS) for bioinformatics support, to the Genomer platform, Biogenouest genomics core facility for its technical support, and the Roscoff Culture Collection (RCC) for culture room facilities. The authors are deeply indebted to the Service Mer & Observation (SMO) of Roscoff, I. Azevero, A. Berthou, L. Brunner, J. Dhinaut, G. Misol, MM. Perrineau, A. Peters, S. Reed, K. Saless, T. Van Berkel, W. Visch, A. Wagner, C. Warwich-Evans for sampling.

## Data availability

All the sequence data (ddRAD-seq) used for this study have been deposited in the European Bioinformatics Institute/European Nucleotide Archive (EBI/ENA) database under the project accessions PRJEB107786 (https://www.ebi.ac.uk/ena/browser/view/PRJEB107786). See Supplementary Table 10 for details.

## Abbreviations

*Â*: allelic richness
AFLP: amplified fragment length polymorphism ANOVA analysis of variance
ATPase: adenosine 5’-TriPhosphatase COI cytochrome-oxydase I
DAPC: discriminant analysis of principal components
ddRAD-seq: double digest restriction-site associated DNA sequencing DEGs differentially expressed genes
EM: Expectation-maximization
eQTL: expression quantitative trait loci
EST: expressed-sequence-tag
FAO: Food and Agriculture Organization
*F*IS: inbreeding coefficient
*F*ST: fixation index (differentiation among populations) GLM generalized linear model
GS: genomic selection
guanosine-5’-triphosphate: 
GWAS: genome-wide association study *H*e expected heterozygosity
HN_4_^+^: ammonium
IMTA: integrated multi-trophic aquaculture LD linkage disequilibrium
MAF: minor allele frequency
MAS: marker-assisted selection
NAD^+^: nicotinamide adenine dinucleotide oxidized form NADH nicotinamide Adenine Dinucleotide
NO_2_: nitrogen dioxide
NO_3_^-^: nitrate
*PÂ*: private allele
PCA: principal component analysis
PES: Provasoli enriched seawater
PVE: phenotypic variance explained
QTL: quantitative trait loci
SLAF-seq: specific locus amplified fragment sequencing SNP single nucleotide polymorphisms
SSR: simple sequence repeats
VCF: variant call format

## Supplementary Table legends

**Supplementary Table 1.** Sampled sites, locations, coordinates and sample sizes of the parental populations.

**Supplementary Table 2.** Number of pseudo-F1 individuals from each population expected in each flask according to the experimental scheme and number observed after the assignment tests.

**Supplementary Table 3.** Self-assignment analysis of the parental sporophytes sampled using 21 SSR loci. n ind.: number of individuals.

**Supplementary Table 4.** SSR loci genotypes of 309 parental *S. latissima* and 494 pseudo-F1 individuals. Number of sample (#), Status (Parental / Pseudo-F1), Name (Sample name with number of flasks for pseudo-F1, i.e. F01 for Flask 01), Site code (site of origin of parental or assigned pseudo-F1 individuals), No. Assig = Not assigned after GeneClass treatment.

**Supplementary Table 5.** Average frequency of null alleles per locus and associated standard deviation. The frequency of null alleles per locus and per population was estimated according to the EM algorithm (Dempster et al. 1977) using FreeNA software (Chapuis and Estoup 2007).

**Supplementary Table 6.** Sample size for each population for morphological and metabolic traits studied in the pseudo-F1.

**Supplementary Table 7.** Morphological and metabolic traits values. The cells that are highlighted in yellow correspond to the parental strains that sired the gametophytes that are in culture at the Roscoff Culture Collection (RCC). For example, 8 gametophytes from the parental stain PTW-02 are cultivated in the RCC collection, they are referred as 12104-12111 in the column "RCC numbers", with the 8 RCC numbers from 12104 to 12111.

**Supplementary Table 8.** Medians for Kruskall-Wallis test comparisons among clusters. For morphological traits: 5 populations in Northern cluster and 7 in Southern cluster. For metabolic traits: 3 populations in Northern cluster and 6 in Southern cluster, except for rhamnose for which only 2 populations were measured in the Northern clade.

**Supplementary Table 9.** GWAS results and annotated genes under the peaks or in a 100kbp window.

**Supplementary Table 10.** Sequence data used in this study.

## Supplementary Figure legends

**Supplementary Figure 1.**
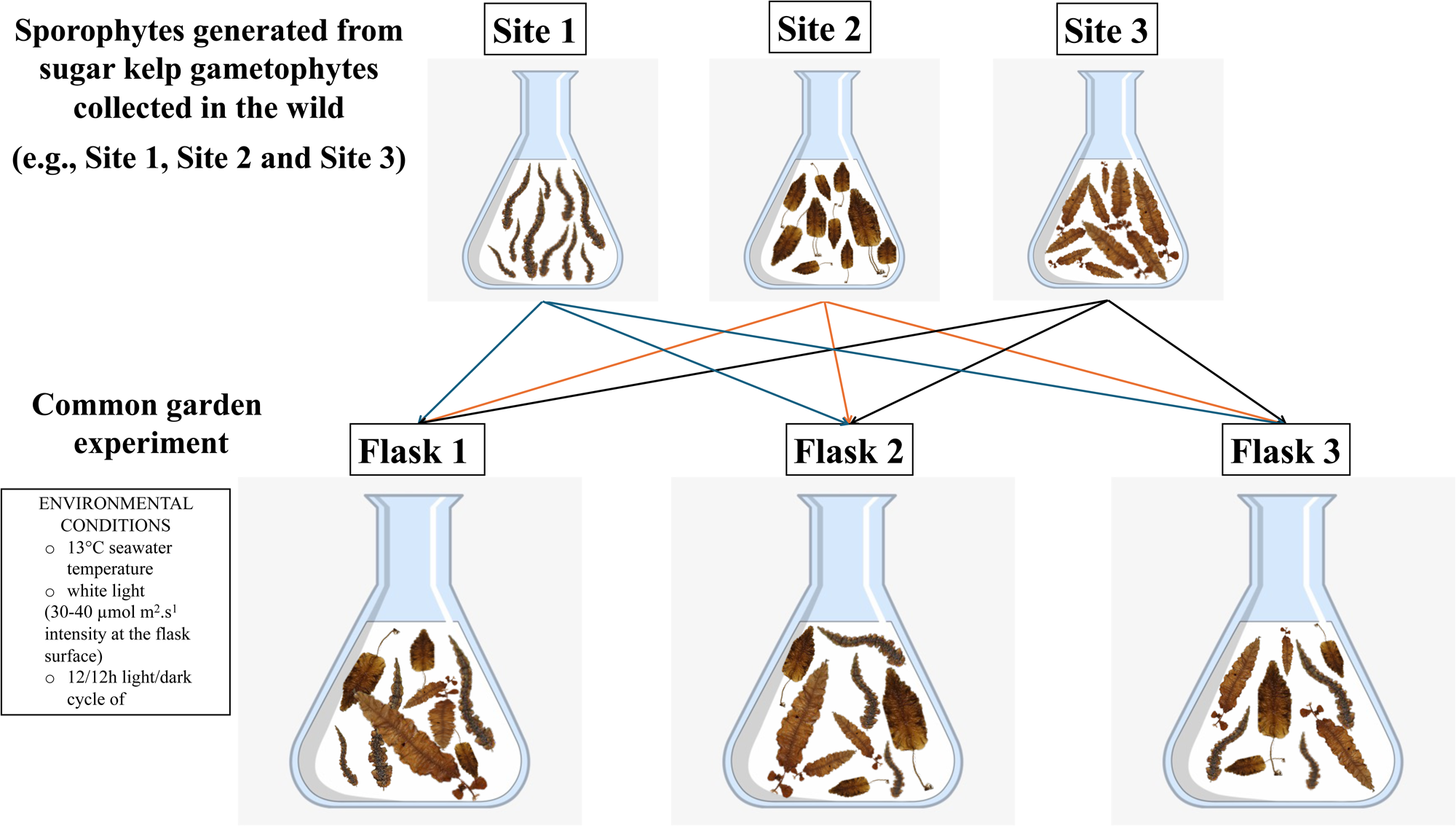
Common garden experiment setup.?

**Supplementary Figure 2.**
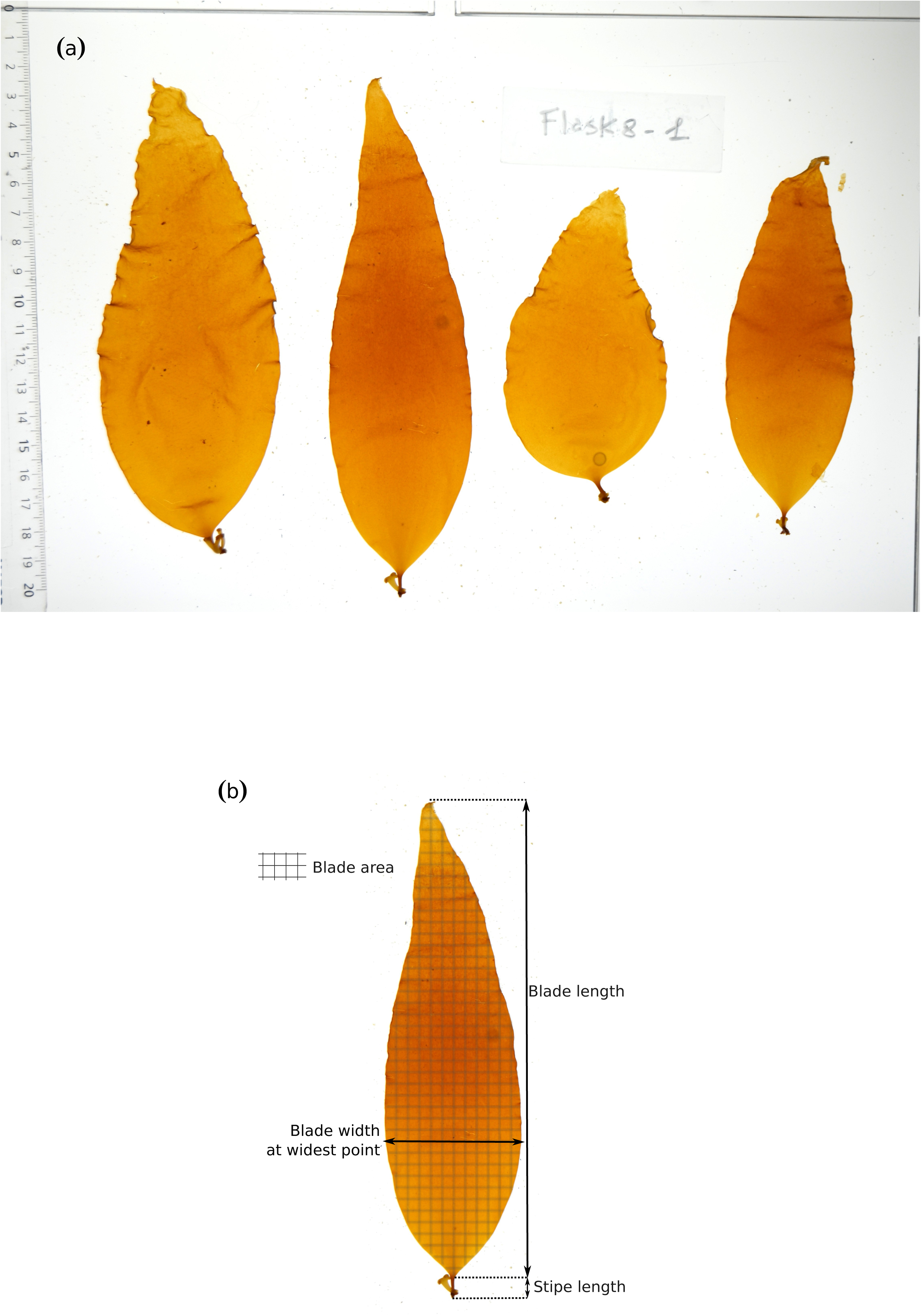
Photo of pseudo-F1 sporophytes analyzed using ImageJ (a) and indication of morphological measurements (b).

**Supplementary Figure 3.**
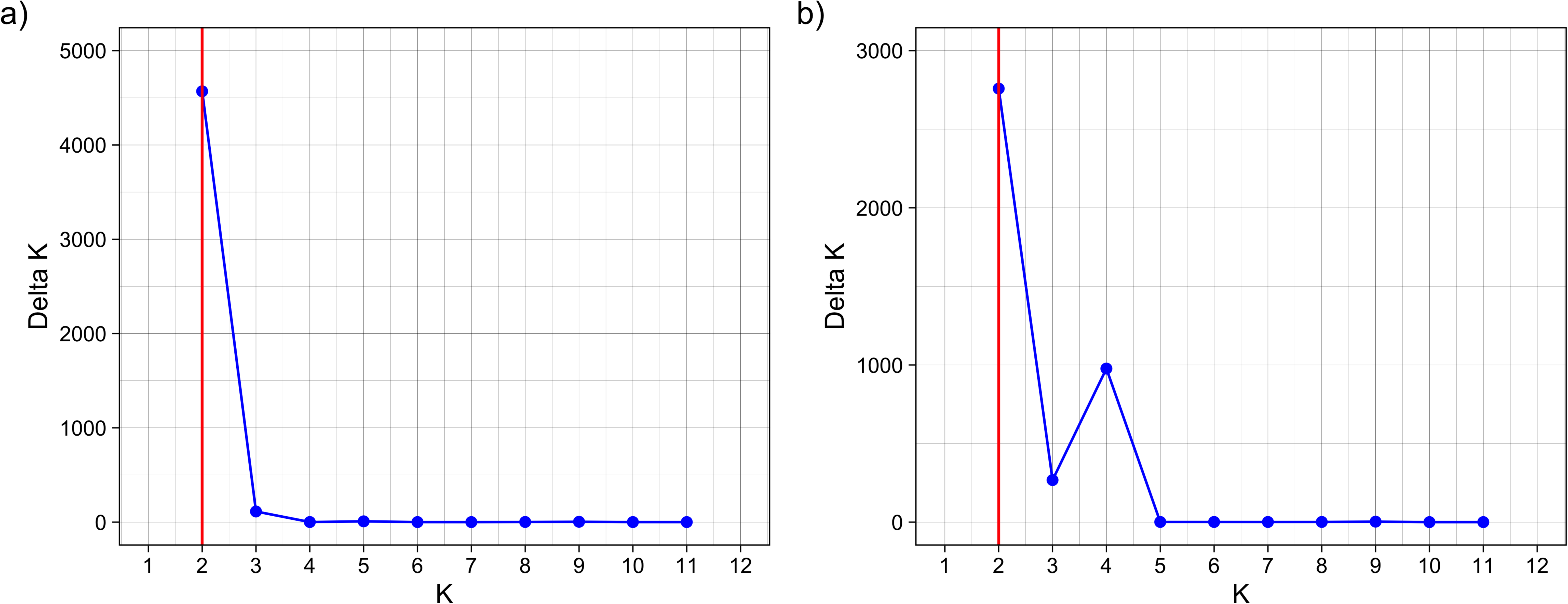
DeltaK values (DeltaK, a measure of the rate of change in the STRUCTURE likelihood function) as a function of K, the number of putative populations. Graphs (a) and (b) show the results for the SSR loci and SNP loci, respectively.

**Supplementary Figure 4.**
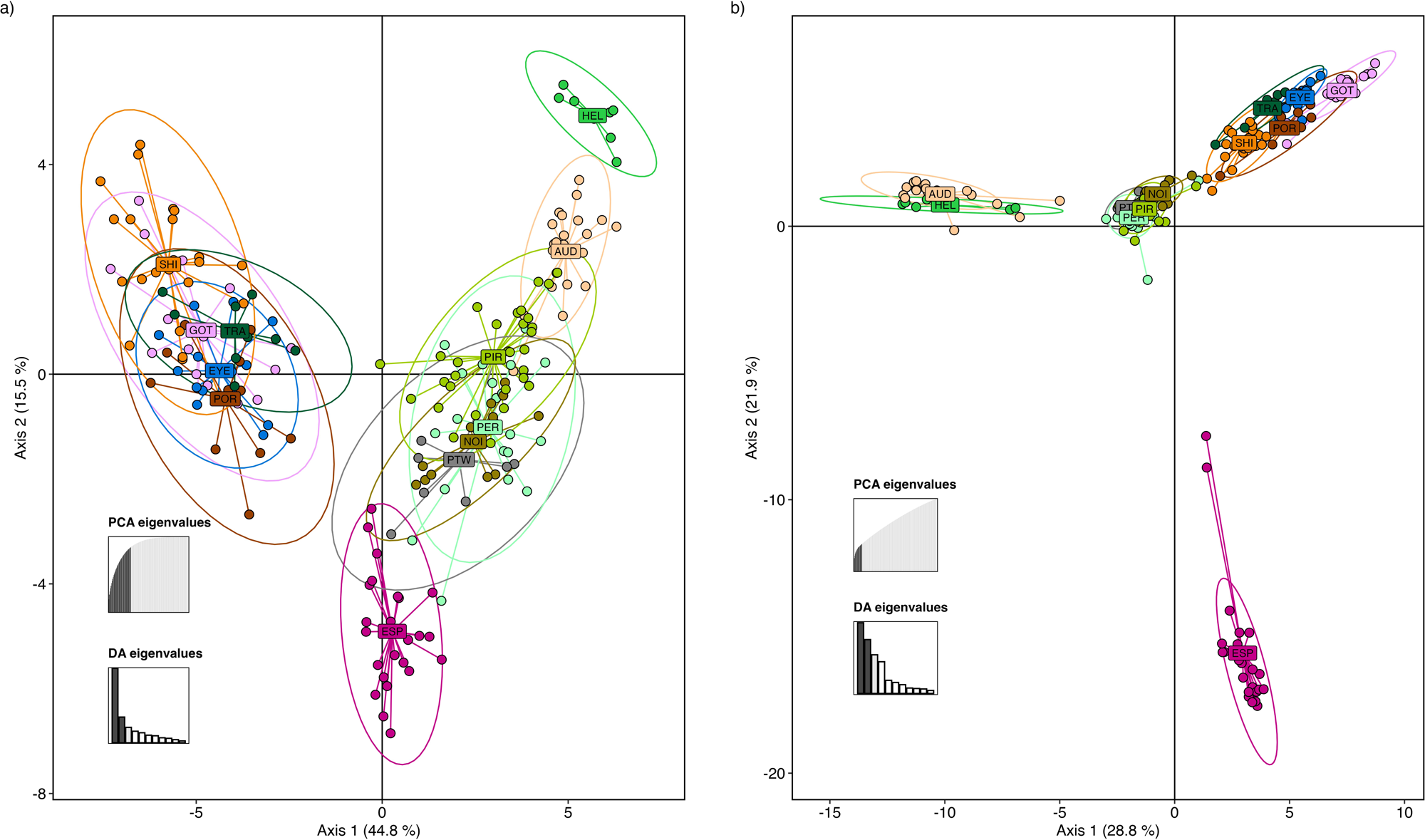
Discriminant Analysis of Principal Components (DAPC) showing the clustering of pseudo-F1 individuals according site level (K=12). Graphs (a) and (b) show the results for the SSR loci and SNP loci, respectively.

**Supplementary Figure 5.**
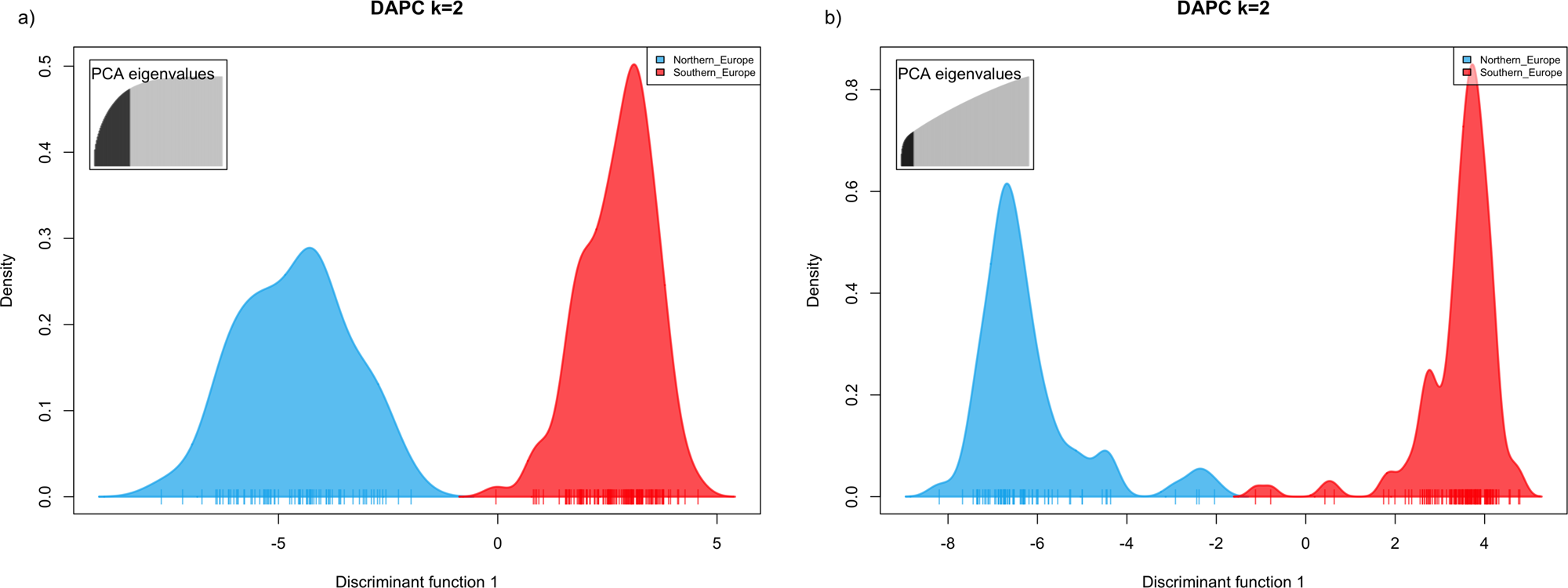
Discriminant Analysis of Principal Components (DAPC) showing the clustering of pseudo-F1 individuals according region level (K=2). Graphs (a) and (b) show the results for the SSR loci and SNP loci, respectively.

**Supplementary Figure 6.**
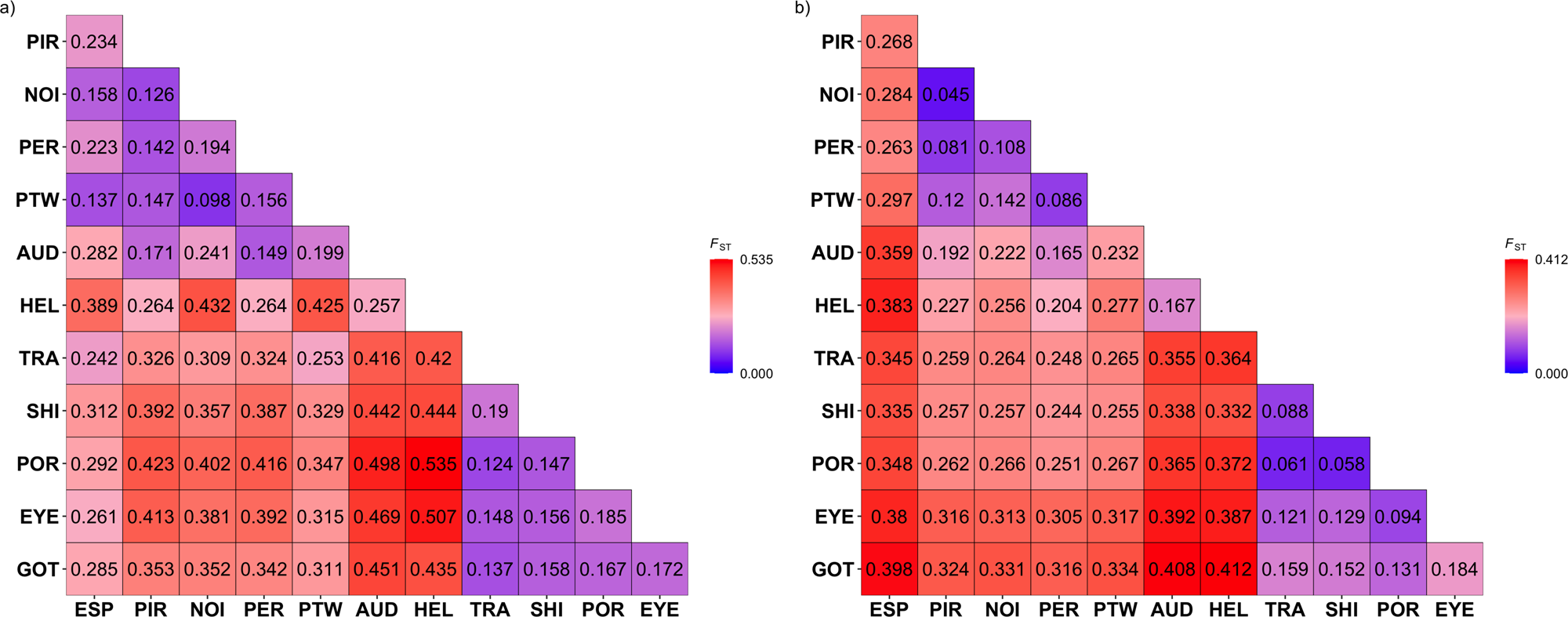
Pairwise genetic differentiation (measured by FST) between 12 *Saccharina latissima* localities and estimated using 21 SSR loci (a) and 71,619 SNPs (b). Pairwise Fst values were significant between all sites.

**Supplementary Figure 7.**
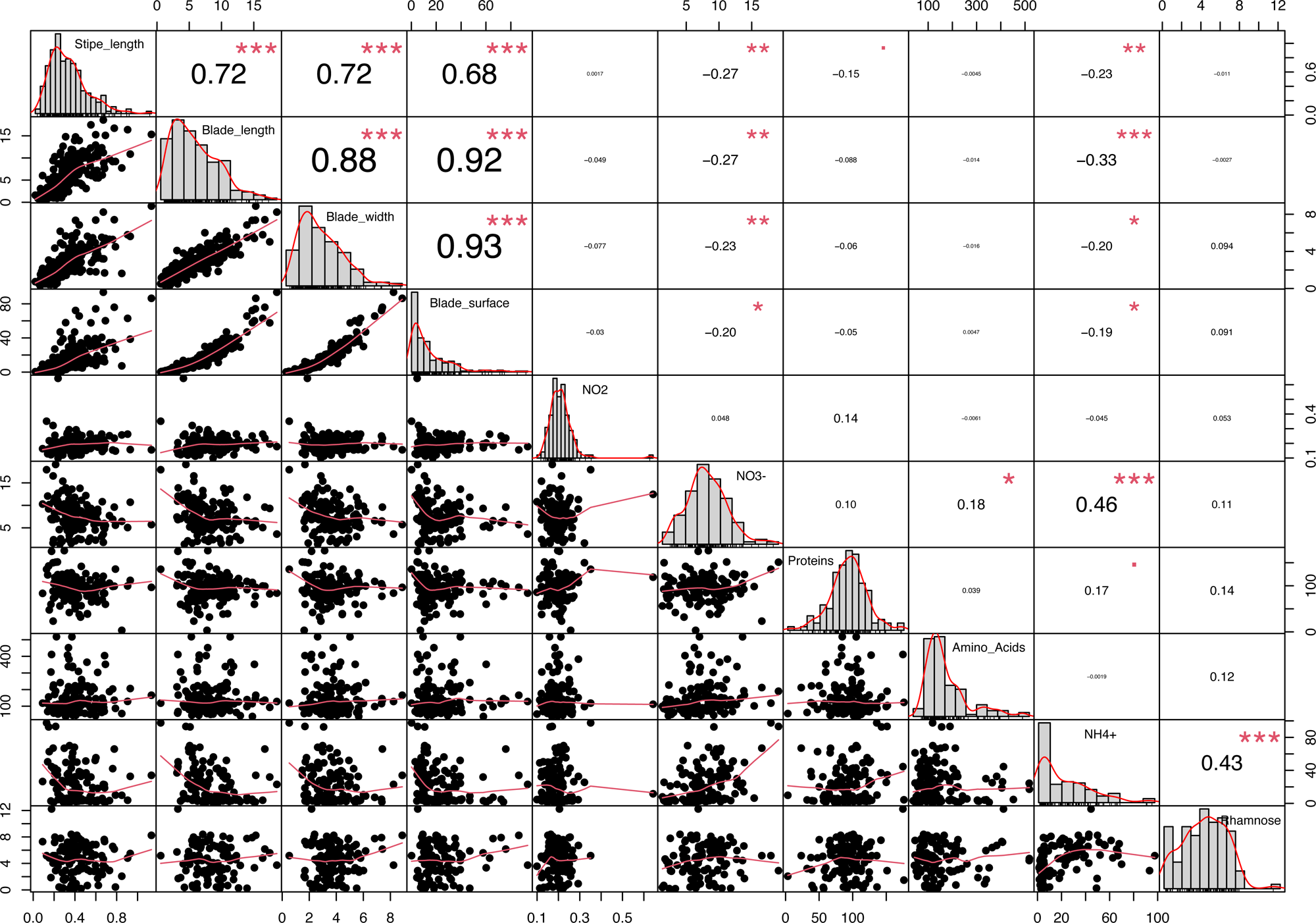
Phenotypic correlation among morphological and metabolic traits analyzed in the pseudo-F1 *S. latissima* population.

**Supplementary Figure 8.**
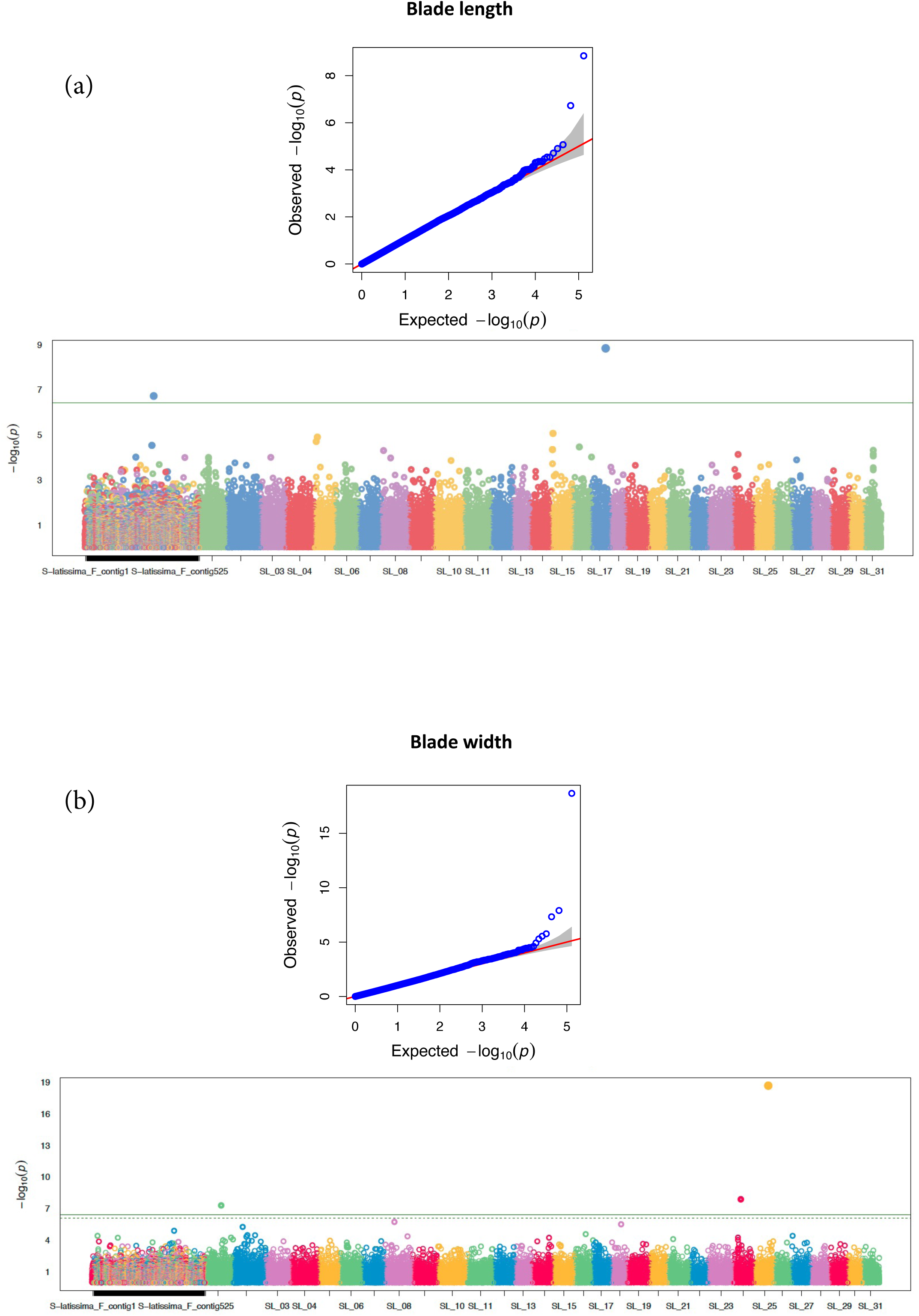
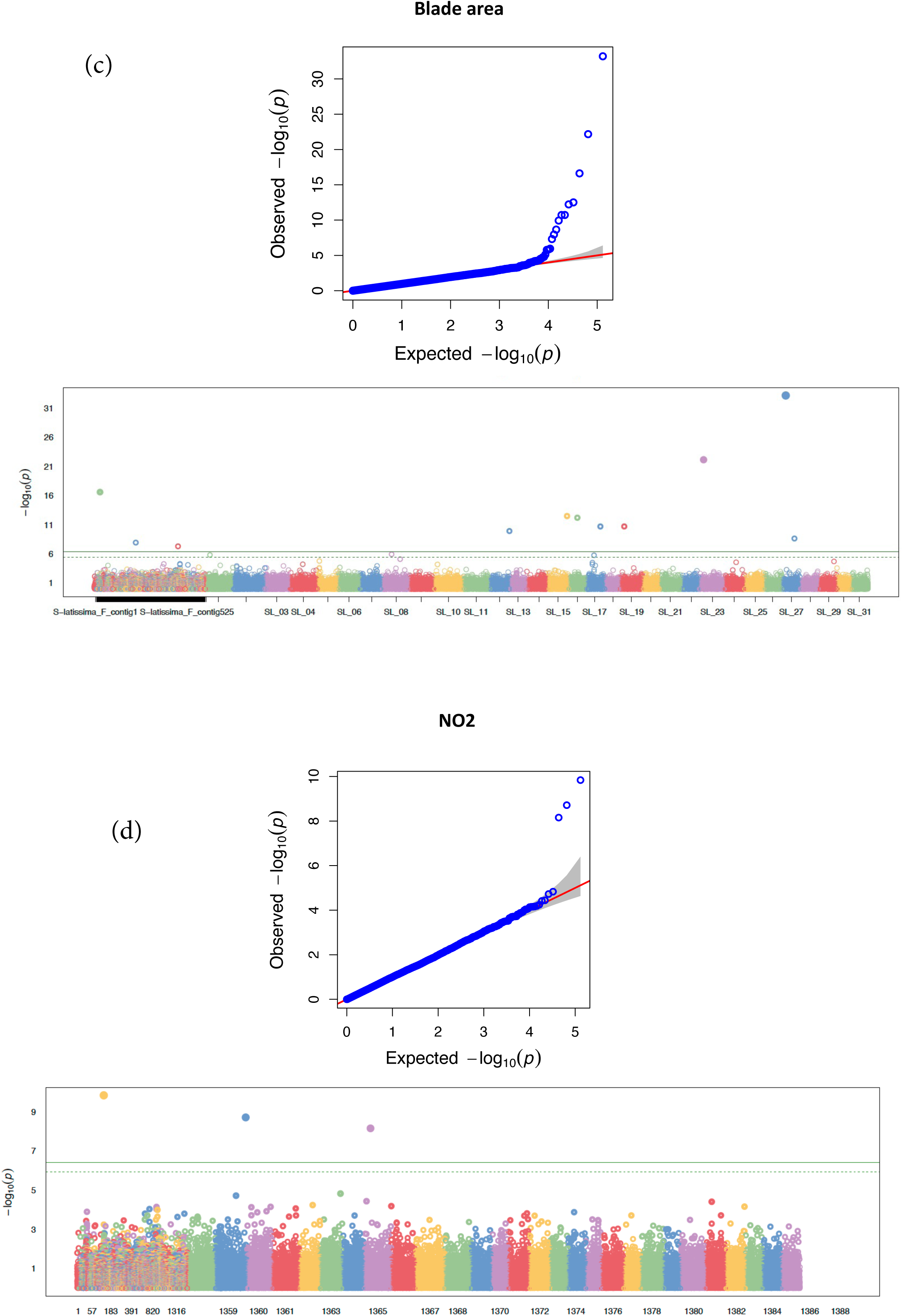
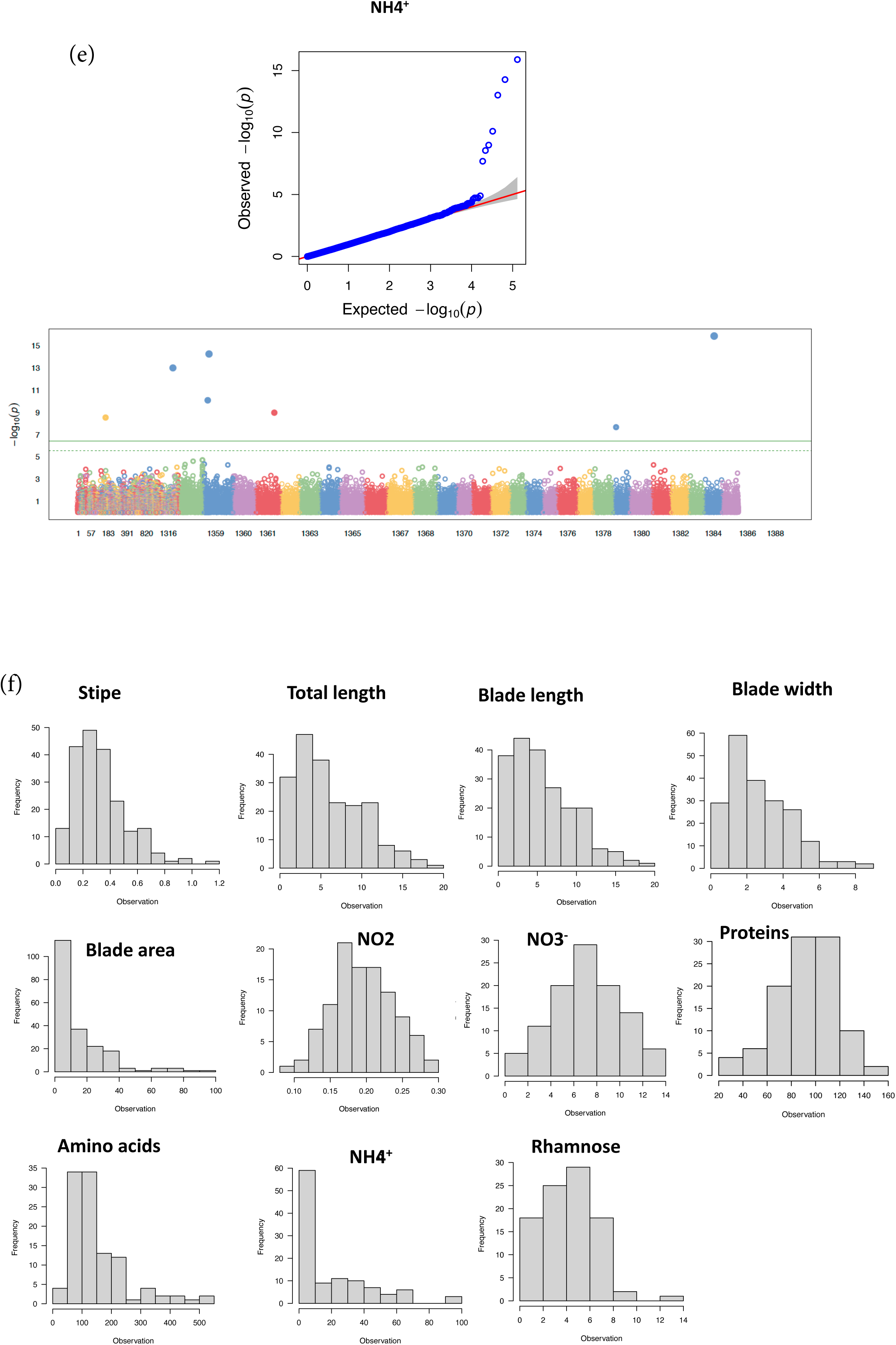
Quantile-quantile (QQ) plot of P-values and Manhattan plot for blade length (a), blade width (b), blade area (c), NO_2_ (d) and NH_4_^+^ (e). Each dot corresponds to a SNP. For QQ plots, the Y-axis is the observed negative base 10 logarithm of the P-values, and the X-axis is the expected observed negative base 10 logarithm of the P-values under the assumption that the P-values follow a uniform distribution. The red line shows the 95% confidence interval for the QQ-plot under the null hypothesis of no association between the SNP and the trait. Manha8an plots summarize GWAS results. The X-axis is the genomic position of each SNP, and the Y-axis is the negative logarithm of the P-value obtained from the GWAS model. Large peaks in the Manha8an plot suggest that the surrounding genomic region has a strong association with the trait. The solid horizontal line corresponds to Bonferroni significance threshold while the dash line corresponds to FDR adjusted significance threshold. Trait distributions are shown in panel f.

**Supplementary Figure 9.**
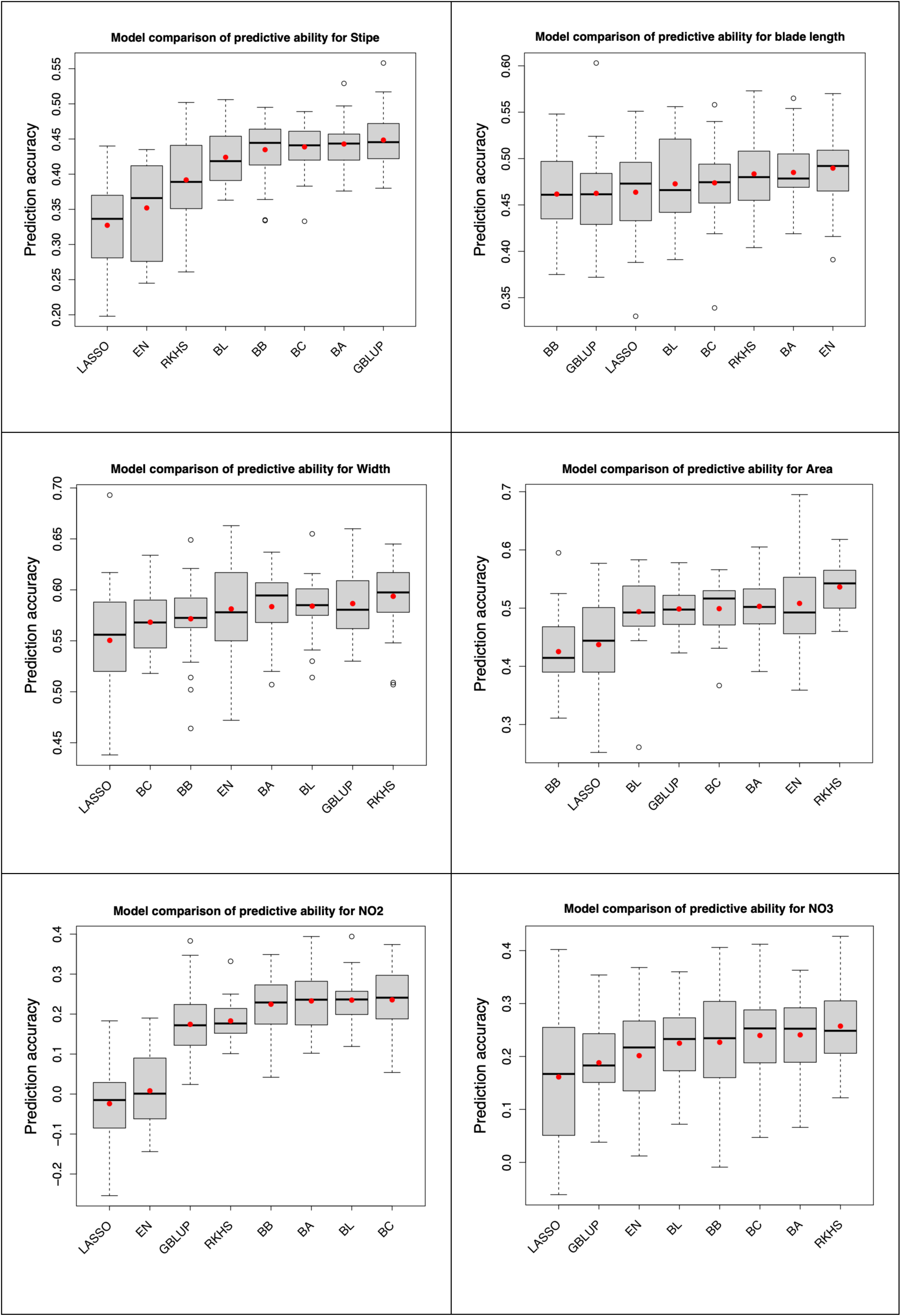
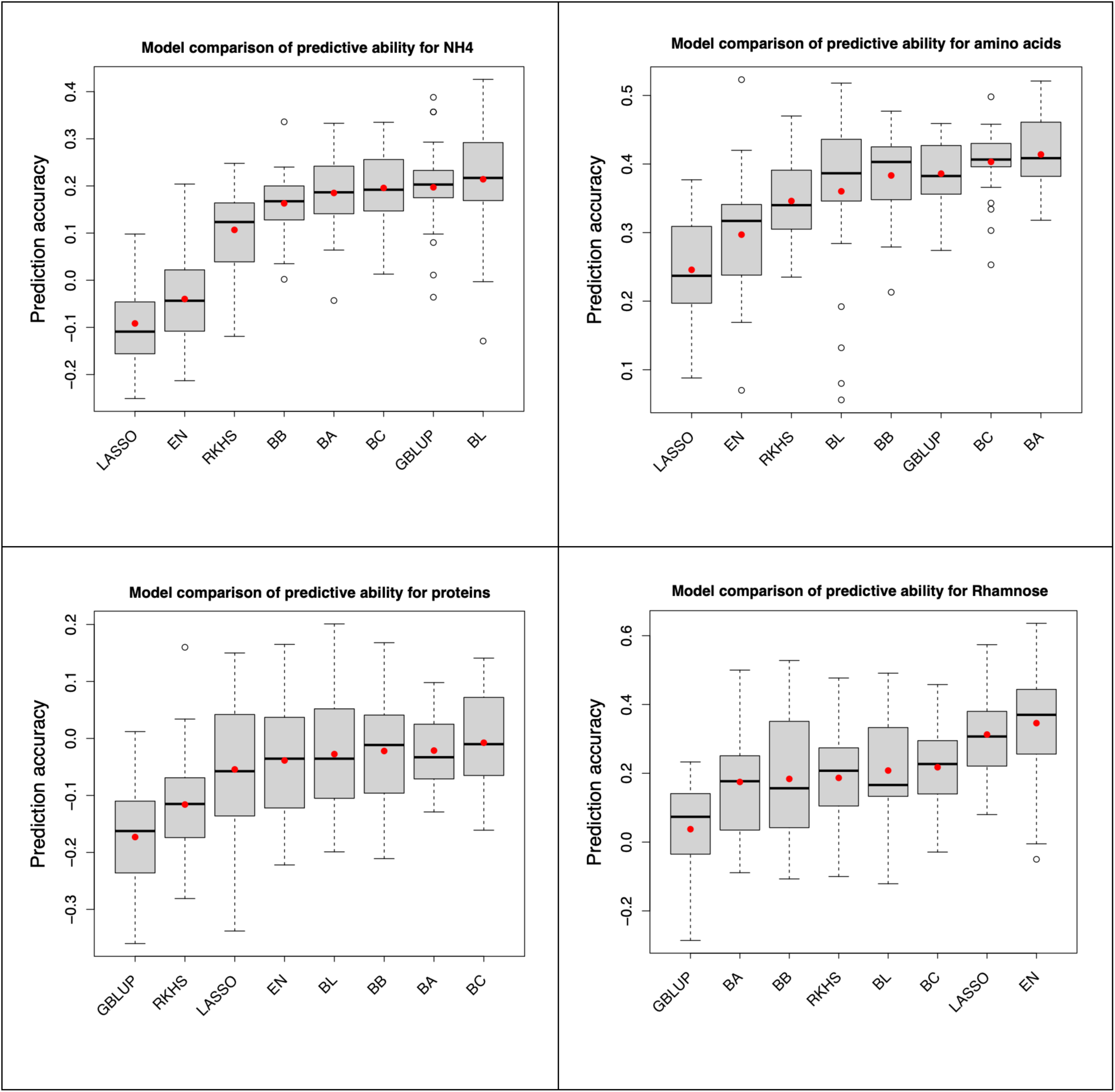
Method comparison for predictive ability (PA). PA is estimated by 5-fold cross-validation and 30 independent replicates. Eight prediction methods were tested: GBLUP, LASSO (Least Absolute Shrinkage and Selection Operator), EN (Elastic Net), RKHS (Reproductive Kernel Hilbert Space), BA (Bayes A), BB (Bayes B), BC (Bayes C) and BL (Bayesian LASSO).

